# Dietary phosphorus restriction enables multigenerational breeding of α-Klotho-deficient rats

**DOI:** 10.64898/2026.10.17.759802

**Authors:** Chihiro Sarai, Yoshikazu Totsuka, Takuro Horii, Sahoko Tamura, Eiko Gouda, Satoko Kojima, Tom Kouki, Nobuhiko Ohno, Atsushi Yoshiki, Tsukasa Ohmori, Hiroshi Kurosu, Hitoshi Endo, Kaoru Tominaga, Izuho Hatada, Takashi Yamamoto, Makoto Kuro-o, Yasumitsu Nagao

**Affiliations:** Center for Experimental Medicine, Jichi Medical University, Shimotsuke, Tochigi, Japan.; Graduate School of Medicine, Jichi Medical University, Shimotsuke, Tochigi, Japan; Faculty of Nutrition, Tokyo Kasei University, Itabashi, Tokyo, Japan.; Institute of Immunology Co., Ltd., Utsunomiya, Tochigi, Japan.; Laboratory of Genome Science, Biosignal Genome Resource Center, Institute for Molecular and Cellular Regulation, Gunma University, Maebashi, Gunma, Japan.; Department of Anatomy, Division of Histology and Cell Biology, Jichi Medical University, Shimotsuke, Tochigi, Japan; Experimental Animal Division, RIKEN BioResource Research Center, Tsukuba, Ibaraki, Japan.; Department of Biochemistry, Jichi Medical University, Shimotsuke, Tochigi, Japan.; Division of Anti-aging Medicine, Center for Molecular Medicine, Jichi Medical University, Shimotsuke, Tochigi, Japan.; Viral Vector Core, Gunma University Initiative for Advanced Research (GIAR), 3-39-22 Showa-machi, Maebashi 371-8511, Gunma, Japan; Genome Editing Innovation Center, Hiroshima University, Higashihiroshima, Hiroshima, Japan.; Division of Mineral Metabolism, Center for Molecular Medicine, Jichi Medical University, Shimotsuke, Tochigi, Japan.

**Author notes:** Correspondence: Yasumitsu Nagao, Center for Experimental Medicine, Jichi Medical University, Shimotsuke, Tochigi, Japan.

## Abstract

Severe α-Klotho deficiency causes early mortality and reproductive impairment, limiting the maintenance and physiological analysis of rat models. We generated rats carrying a CRISPR/Cas9-mediated α-Klotho deletion spanning exons 1–5 and evaluated dietary phosphorus restriction primarily in the Wistar-Imamichi background. Maternal low-phosphorus (LP) feeding from mating through lactation was associated with higher overall survival to weaning (92.6% versus 85.8% under normal-phosphorus conditions) and a higher proportion of KO animals among weaned survivors (34/113 versus 23/145). Maternal LP feeding also reduced plasma phosphorus in KO pups. Continued LP feeding after weaning supported growth, gonadal development, and natural KO × KO breeding. Across 15 litters, 74 of 85 offspring survived to weaning, and KO colonies were maintained to at least the fifth generation. We additionally characterized 24-h metabolic rhythms in LP-maintained KO animals using indirect calorimetry. These profiles remained heterogeneous under a light–dark cycle. In conclusion, dietary phosphorus restriction provides a reliable strategy for the multigenerational maintenance of severe α-Klotho-deficient rats, establishing a maintainable model to investigate residual metabolic and physiological phenotypes.

## Introduction

α-Klotho functions as an essential co-receptor for fibroblast growth factor 23 (FGF23) and plays a central role in regulating renal phosphate excretion and vitamin D homeostasis (Shimada et al., 2004; Kurosu et al., 2006; Urakawa et al., 2006; Nakatani et al., 2009; Hu et al., 2010). Loss of α-Klotho results in hyperphosphatemia, systemic ectopic calcification, infertility, and markedly shortened lifespan, a phenotype often described as resembling accelerated aging (Kuro-o et al., 1997; Ohnishi and Razzaque, 2010; John et al., 2011).

Previous studies have demonstrated phenotypic improvement in hypomorphic Klotho-mutant mice through dietary intervention and restoration of reproductive ability in Klotho/NaPi2a double-knockout mice through genetic reduction of phosphate burden (Morishita et al., 2001; Ohnishi and Razzaque, 2010). These findings support the contribution of phosphate toxicity to disease severity in α-Klotho-deficient states (Kuro-o, 2010; Razzaque, 2011). The originally reported Klotho-mutant mouse carries a severe hypomorphic allele with residual expression (Kuro-o et al., 1997; Kuro-o, 2009), whereas knockout models have also been studied. We therefore focused on establishing dietary conditions that permit natural breeding and multigenerational maintenance of rats carrying a large α-Klotho deletion.

Early mortality and reproductive impairment limit the maintenance of severe α-Klotho-deficient rat lines, despite the practical advantages of rats for longitudinal physiological studies. Their larger body size facilitates repeated sampling, surgical procedures, and detailed assessment of mineral metabolism (Jacob, 2010; Szpirer, 2020; Chenouard et al., 2021). Genome editing has enabled efficient generation of genetically modified rats (Shao et al., 2014; Yoshimi et al., 2014; Sato et al., 2022).

However, early lethality and infertility necessitate husbandry conditions that permit long-term maintenance and physiological analysis of mutant rats.

We tested whether LP feeding from mating through lactation, followed by continued LP feeding after weaning, could support offspring recovery and natural KO × KO breeding across generations. We focused on the Wistar-Imamichi (WI) background because of its stable 4-day estrous cyclicity and reliable breeding performance. We assessed overall survival and genotype distribution at weaning, plasma mineral parameters, growth, gonadal development, and natural breeding. Estrous cyclicity and breeding outcomes were evaluated as indicators of reproductive function. We additionally characterized 24-h metabolic rhythms to assess residual physiological differences in LP-maintained KO animals.

## Results

### Generation and sequence confirmation of rats carrying a large α-Klotho deletion

We generated an approximately 40-kb deletion encompassing the proximal promoter region and exons 1–5 of the rat α-Klotho gene using CRISPR/Cas9 (Fig. 1a). Genotyping combined 5′ wild-type allele PCR, 3′ wild-type allele PCR, and deletion-junction PCR to distinguish wild-type (WT), heterozygous (Het), and knockout (KO) animals. Sanger sequencing confirmed formation of the intended deletion junction (Fig. 1b).

**Figure 1.**
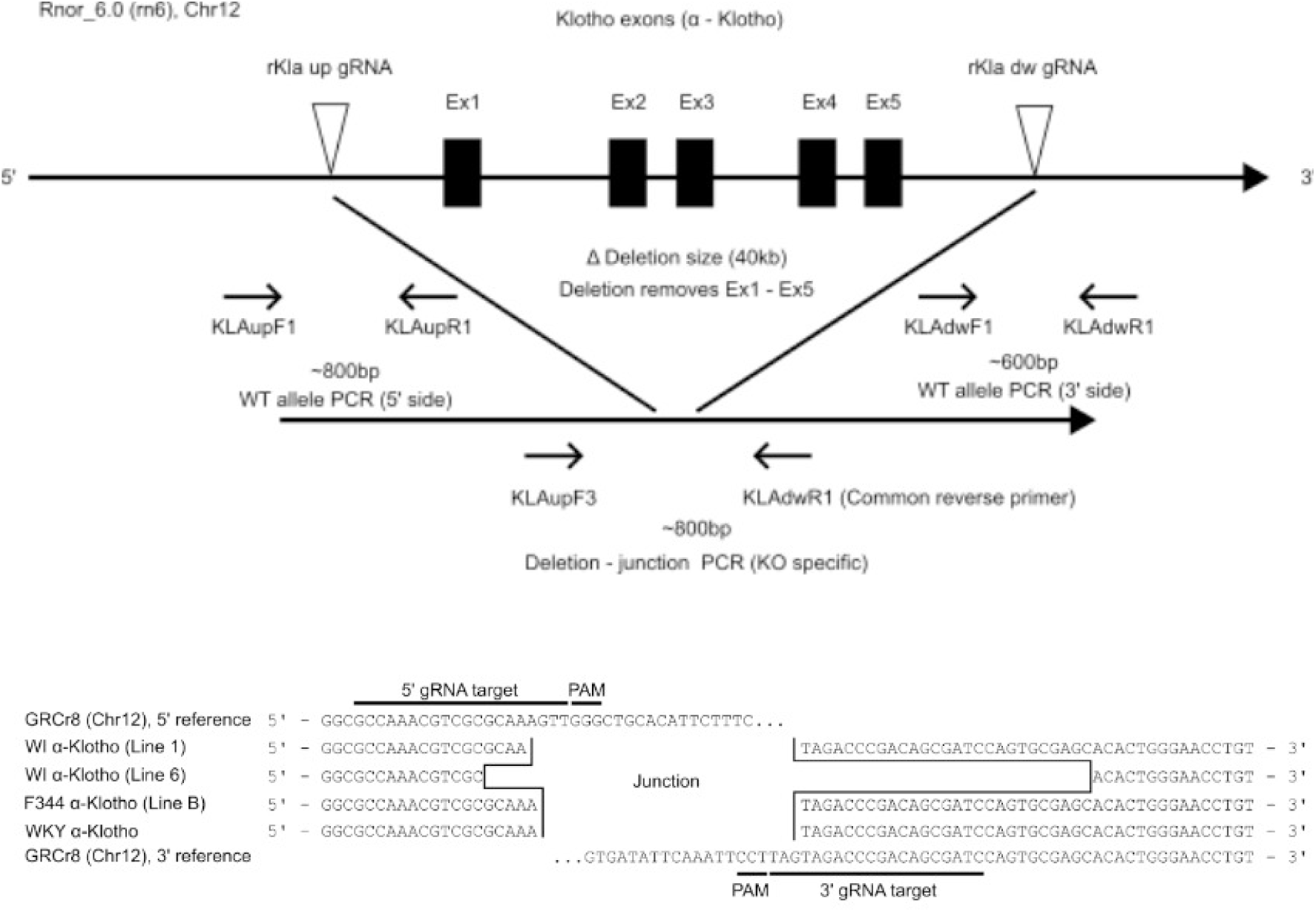
Generation and sequence confirmation of rats carrying a large α-Klotho deletion. a, CRISPR/Cas9 design for deletion of the proximal promoter region and exons 1–5 of the rat α-Klotho gene (approximately 40 kb). Guide RNAs target the 5′ and 3′ flanking regions. Genotyping used 5′ wild-type allele, 3′ wild-type allele, and deletion-junction PCR assays. WT animals yielded wild-type allele bands only, Het animals yielded both wild-type and deletion-junction bands, and KO animals yielded the deletion-junction band only. b, Sequence confirmation of deletion-junction alleles in rat lines used for breeding. Nine WI, two F344, and one WKY line were sequence-confirmed; two WI, one F344, and one WKY line were used for breeding observations. WI, Wistar-Imamichi; F344, Fischer 344; WKY, Wistar-Kyoto.

Sequence confirmation was performed for nine independently derived WI lines, and two of these lines were used for breeding experiments. WI lines derived from two independent founders were maintained. Additional rat backgrounds and mouse lines are described in the Supplementary Results.

### Maternal LP feeding was associated with higher overall survival and KO recovery at weaning

We evaluated overall survival to weaning and the genotype distribution among weaned offspring under NP and maternal LP conditions (Table 1). Under NP conditions, WI heterozygous intercrosses produced 169 pups from 14 litters, with a mean litter size of 12.1 ± 2.9. Of these, 145 survived to weaning (85.8%). Their genotype distribution was WT:Het:KO = 58:64:23, corresponding to 40.0%, 44.1%, and 15.9%, respectively. This distribution differed from the expected Mendelian ratio of 1:2:1 (χ² = 18.89, P = 7.91 × 10⁻⁵).

**Table 1.** Overall survival and genotype distribution at weaning in WI heterozygous intercrosses.

| Strain | Diet | Litters | Born | Weaned | Overall survival to weaning (%) | Died | WT | Het | KO | $\chi^2$ | P value |
| --- | --- | --- | --- | --- | --- | --- | --- | --- | --- | --- | --- |
| WI α-Klotho | NP | 14 | 169 | 145 | 85.8 | 24 | 58 (40.0%) | 64 (44.1%) | 23 (15.9%) | 18.89 | $7.91 \times 10^{-5}$ |
| WI α-Klotho | LP | 10 | 122 | 113 | 92.6 | 9 | 36 (31.9%) | 43 (38.1%) | 34 (30.1%) | 6.52 | 0.0383 |

Breeding outcomes under NP and LP conditions in WI rats. Overall survival to weaning was calculated as Weaned / Born × 100. Weaning survival refers to pups surviving to postnatal days 20–32. WT, Het, and KO counts are shown as n (%), with percentages calculated among all weaned survivors in each dietary group. Genotypes were available for all survivors. Deviation from the expected Mendelian 1:2:1 ratio was assessed using the genotype counts by chi-square goodness-of-fit testing. These tests do not directly compare NP and LP groups. Mortality before genotyping prevents determination of complete genotype-specific survival. NP, normal-phosphorus diet; LP, low-phosphorus diet; WI, Wistar-Imamichi; WT, wild type; Het, heterozygous; KO, knockout.

When females were switched to LP feeding at the start of mating and maintained on LP throughout gestation and lactation, 122 pups were obtained from 10 litters, with a mean litter size of 12.2 ± 3.3. Of these, 113 survived to weaning (92.6%). The genotype distribution among survivors was WT:Het:KO = 36:43:34, corresponding to 31.9%, 38.1%, and 30.1%, respectively. The distribution still differed from the expected 1:2:1 ratio (χ² = 6.52, P = 0.0383).

Maternal LP feeding was thus associated with improved overall survival to weaning and a markedly increased yield of viable KO pups.

### Maternal LP feeding reduced plasma phosphorus and altered calcium and magnesium in KO pups

Plasma phosphorus (P), calcium (Ca), and magnesium (Mg) were measured in pups at postnatal days 20–32 (Fig. 2). Under NP conditions, the median plasma P concentration in KO pups was 13.65 mg/dL. Under maternal LP conditions, it was reduced to 8.65 mg/dL (Holm-adjusted P = 0.0051). Plasma Ca increased from 10.70 to 12.45 mg/dL (Holm-adjusted P = 0.0196), whereas plasma Mg decreased from 2.70 to 2.05 mg/dL (Holm-adjusted P = 0.0287).

**Figure 2.**
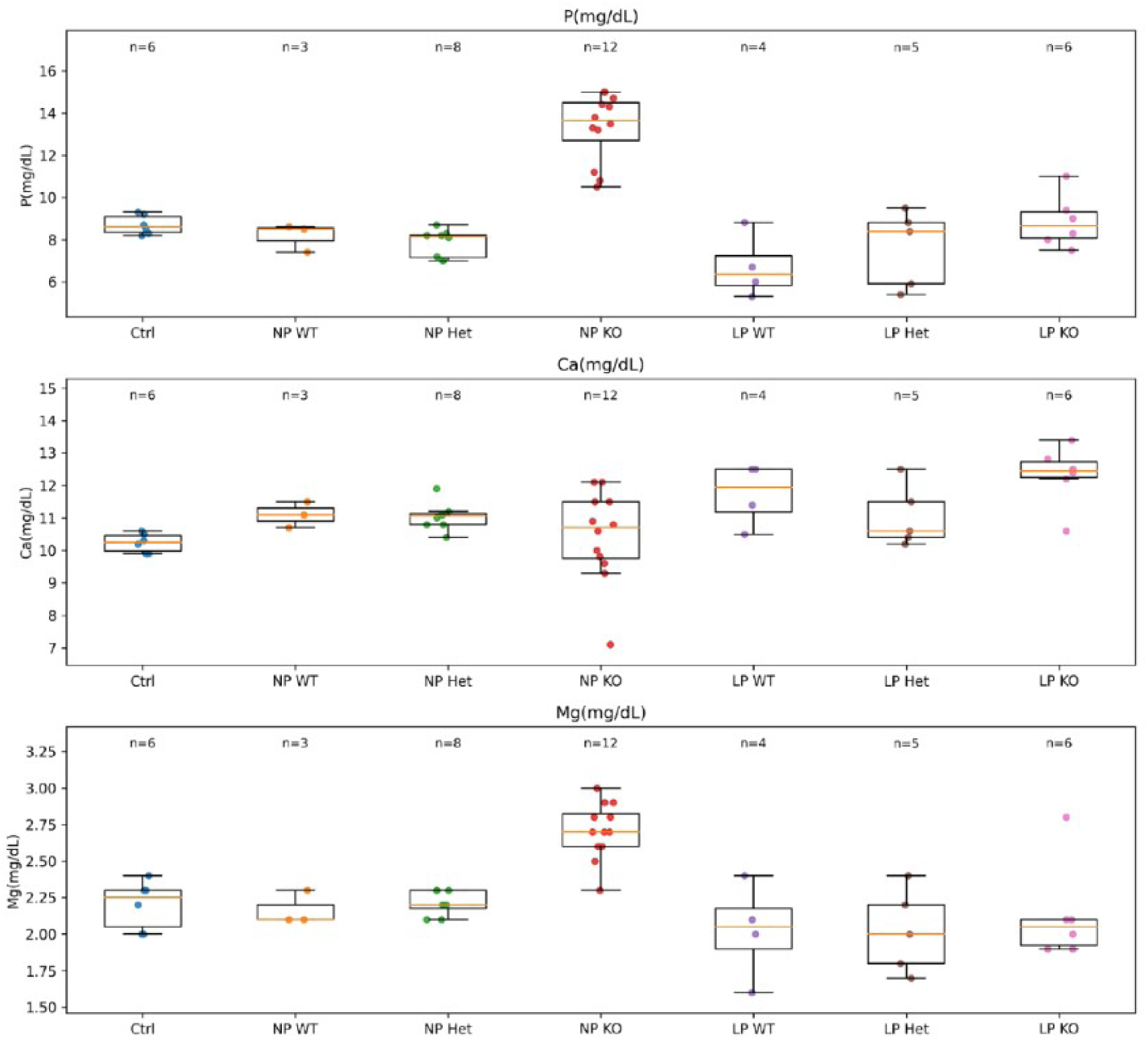
Plasma mineral concentrations in pups under maternal NP and LP conditions. Plasma phosphorus (P), calcium (Ca), and magnesium (Mg) in pups sampled at postnatal days 20–32 under NP and maternal LP conditions. Individual values and summary statistics are shown. NP–LP comparisons within each genotype used two-sided Mann–Whitney U tests with Holm correction for multiple comparisons. Significant differences are indicated. The data describe mineral concentrations under the tested maternal dietary conditions and do not demonstrate complete normalization of mineral homeostasis.

These changes were relatively modest in WT and Het pups. Thus, maternal LP feeding effectively lowered plasma phosphorus in KO pups, accompanied by reciprocal shifts in calcium and magnesium levels.

### Body weight differences at P21 were attenuated under LP conditions

We examined body weight at P21 under NP and LP conditions. To ensure age-matched comparisons, the main analysis was restricted to WI animals evaluated at P21 (Fig. 3). Under LP conditions, the severe growth retardation characteristic of KO pups was substantially attenuated at P21.

**Figure 3.**
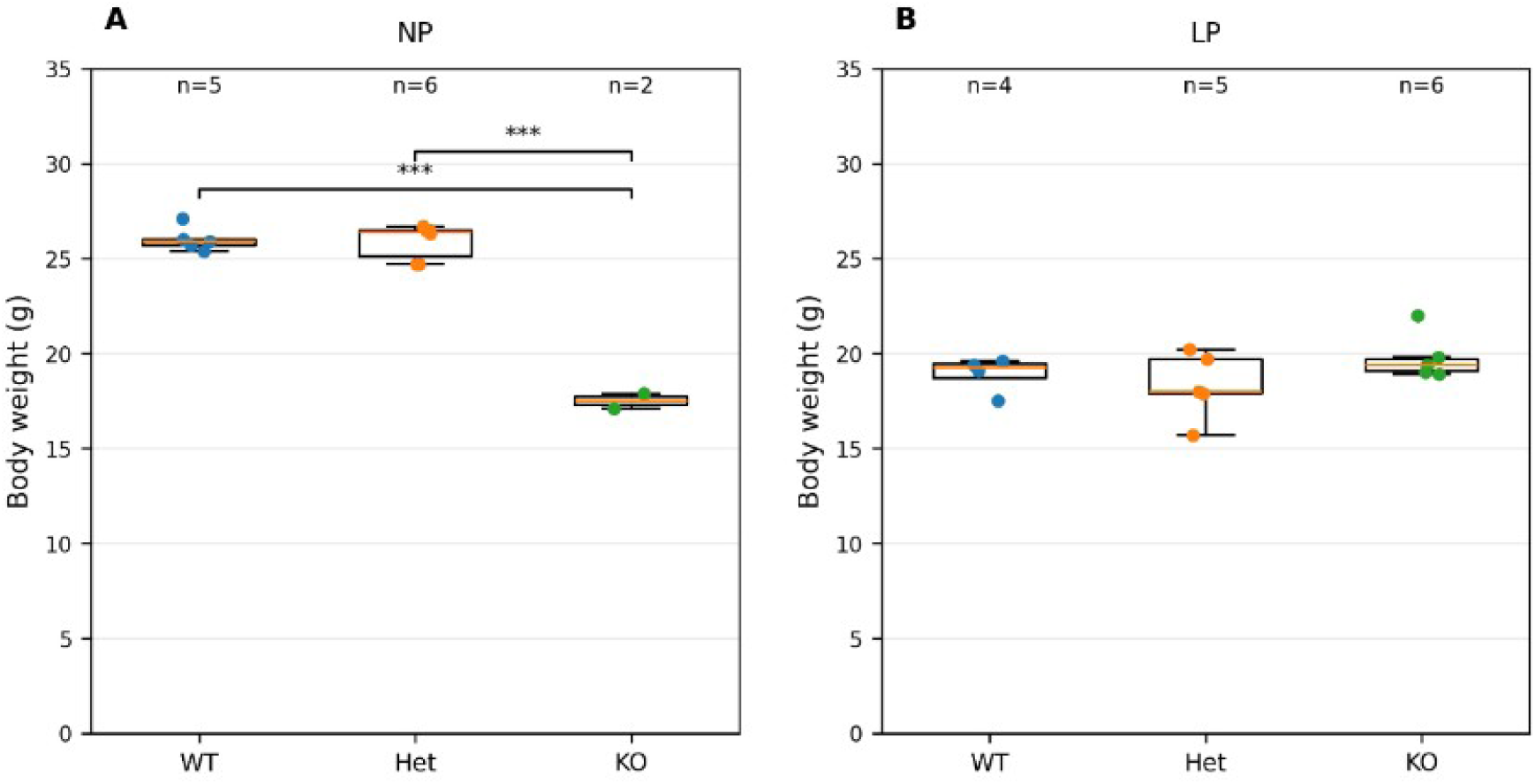
Body weight of WI rats at P21 under NP and LP conditions. Body weight of WT, Het, and KO WI rats at postnatal day 21 (P21). Each point represents one animal. Boxes show the median and interquartile range; whiskers extend to the most extreme observations within 1.5 × IQR. All individual observations are plotted. Group sizes are indicated above each group. Two-way ANOVA was followed by Tukey’s multiple-comparisons test. In the NP panel, WT–KO and Het–KO comparisons were significant (***P < 0.001). These age-matched observations do not isolate the effect of post-weaning intervention.

### Continued LP feeding supported reproductive development and multigenerational breeding

We next examined whether continued phosphorus restriction after weaning affected indices associated with reproductive maturation. Under NP conditions, KO rats remained markedly underweight, and both testis weight in males and ovary weight in females were low. By contrast, KO animals maintained on LP after weaning exhibited greater body weight together with increased kidney and gonadal weights, consistent with improved organ growth (Figs. 4 and 5).

**Figure 4.**
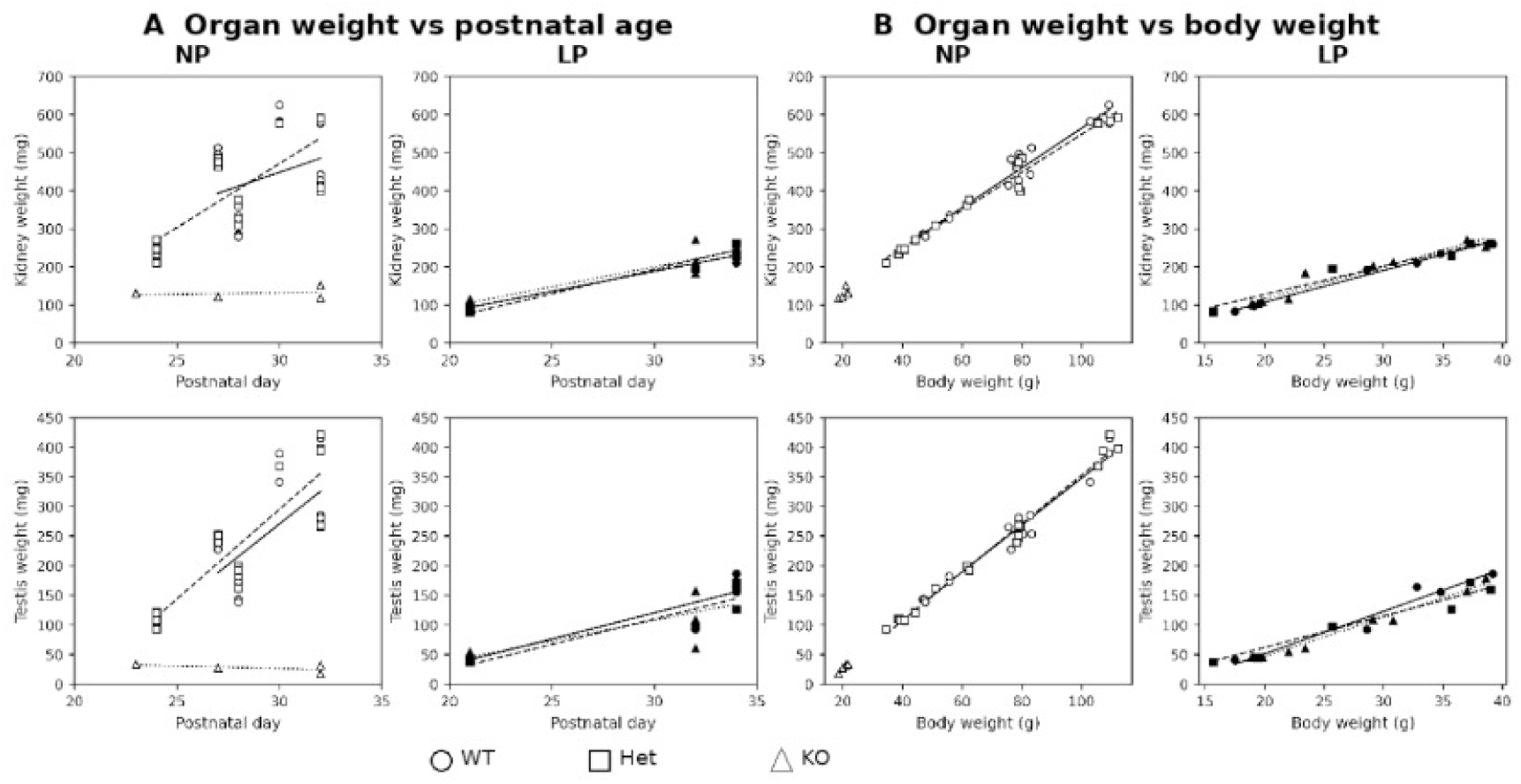
Kidney and testis growth in male rats under NP and LP conditions. a, Relationships between postnatal age and organ weight. b, Relationships between body weight and organ weight. Upper panels show kidney weight and lower panels show testis weight. Panels are separated by NP and LP conditions. Each point represents one animal. Circles, squares, and triangles denote WT, Het, and KO animals, respectively. Open and filled symbols denote NP and LP groups, respectively. Solid, dashed, and dotted lines denote WT, Het, and KO trends, respectively; regression lines are shown for visualization only. LP-maintained KO animals showed higher body and organ weight ranges than NP-maintained KO animals.

**Figure 5.**
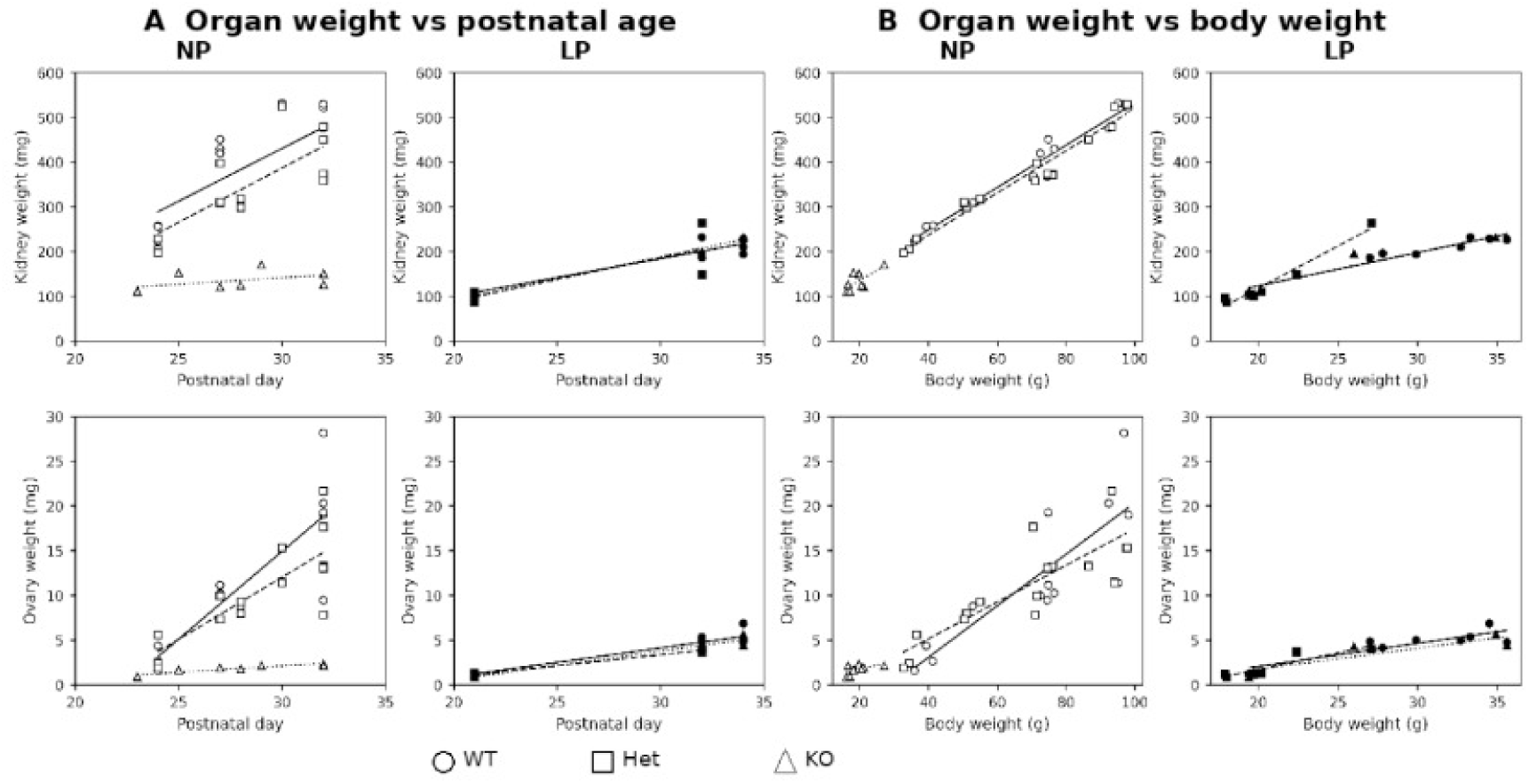
Kidney and ovary growth in female rats under NP and LP conditions. a, Relationships between postnatal age and organ weight. b, Relationships between body weight and organ weight. Upper panels show kidney weight and lower panels show ovary weight. Panels are separated by NP and LP conditions. Each point represents one animal. Circles, squares, and triangles denote WT, Het, and KO animals, respectively. Open and filled symbols denote NP and LP groups, respectively. Solid, dashed, and dotted lines denote WT, Het, and KO trends, respectively; regression lines are shown for visualization only. LP-maintained KO animals showed higher body and organ weight ranges than NP-maintained KO animals.

In male KO animals, NP conditions were associated with confinement to a low-body-weight range and reduced kidney and testis weights, whereas LP conditions were associated with a shift toward a higher body-weight range together with increased kidney and testis weights (Fig. 4). A similar pattern was observed in females: ovary weights remained low in KO animals under NP conditions but were increased in LP-maintained KO animals in parallel with body weight gain, and kidney weights were likewise higher under LP conditions than under NP conditions (Fig. 5).

Histological analysis further showed improvement of testicular architecture in LP-maintained KO males (Fig. 6A–D). Spermatozoa were also observed in the seminiferous tubule lumen of a 10-week-old KO male maintained under LP conditions (Fig. 6E). Consistently, scanning electron microscopy of sperm collected from rescued KO males did not reveal obvious morphological abnormalities within the range examined (Fig. 7).

**Figure 6.**
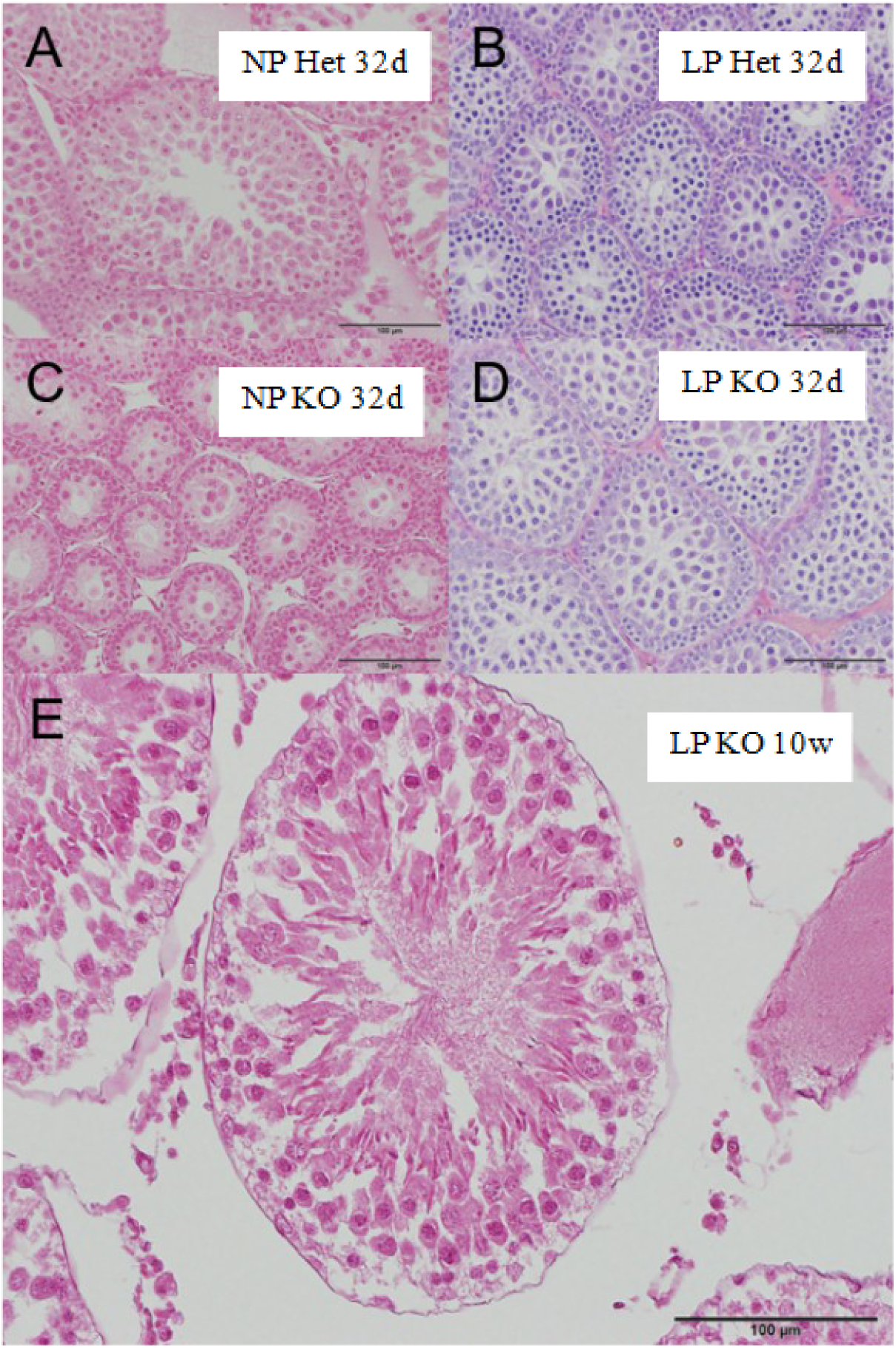
Testicular histology in KO rats under NP and LP conditions. Representative hematoxylin and eosin (H&E)-stained testis sections. A, NP Het; B, LP Het; C, NP KO; D, LP KO. A–D show testes collected at P32 and were acquired at the same magnification. E, Testis from a 10-week-old KO male maintained under LP conditions, showing spermatozoa in the seminiferous tubule lumen. Scale bars, 100 μm.

**Figure 7.**
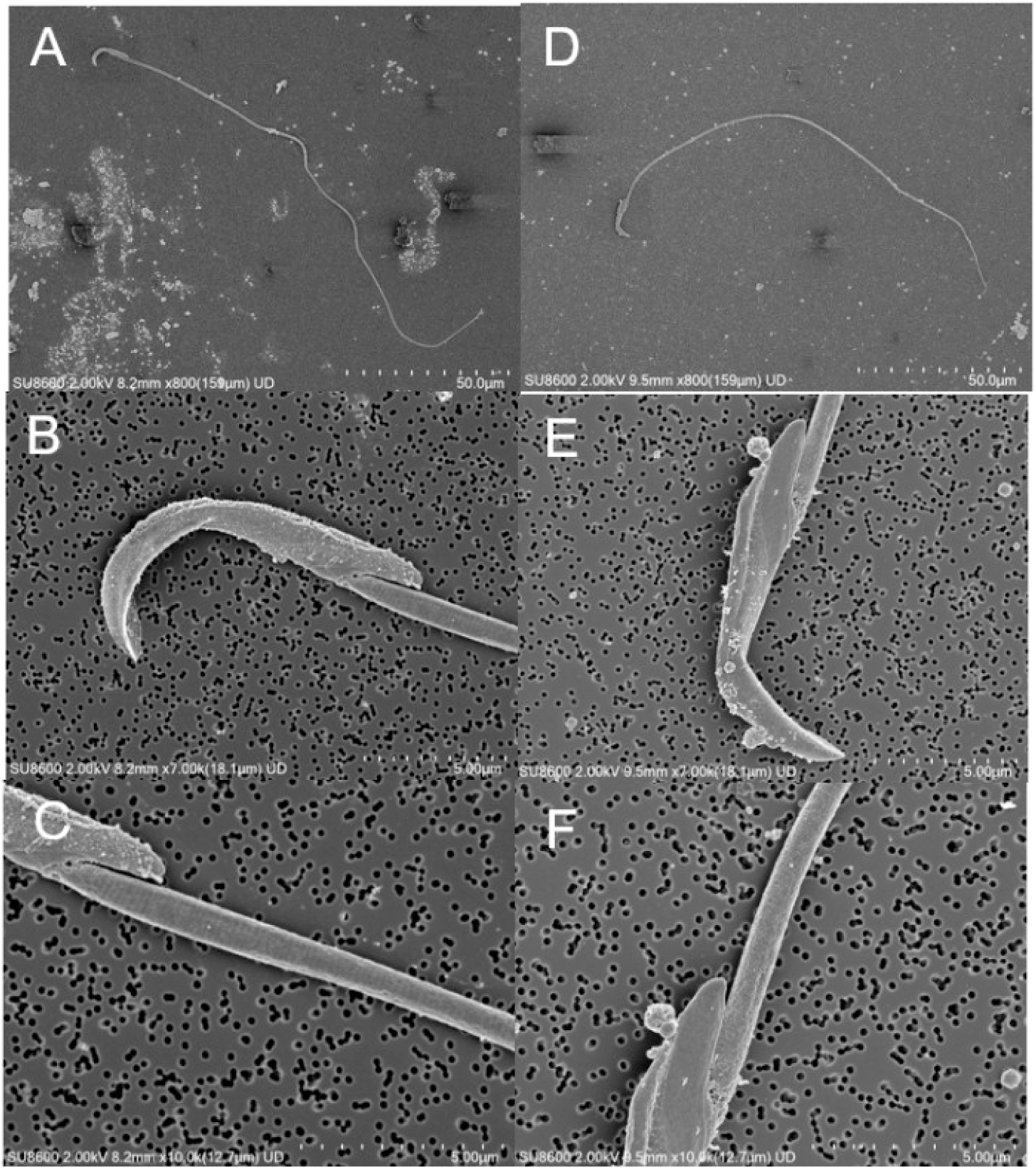
Sperm morphology in WT controls and LP-maintained KO males. Representative scanning electron microscopy (SEM) images of spermatozoa from NP-maintained WT rats (A–C) and LP-maintained KO rats (D–F). (A, D) Whole spermatozoa. (B, E) Views of the sperm head. (C, F) Views of the midpiece. Scale bars, 50 µm (A, D) and 5 µm (B, C, E, F).

Natural mating between WI KO animals maintained under LP conditions produced 85 pups from 15 litters, of which 74 (87.1%) survived to weaning. These outcomes enabled maintenance of the KO colony to at least the fifth generation (F5) (Fig. 8, Table 2).

**Figure 8.**
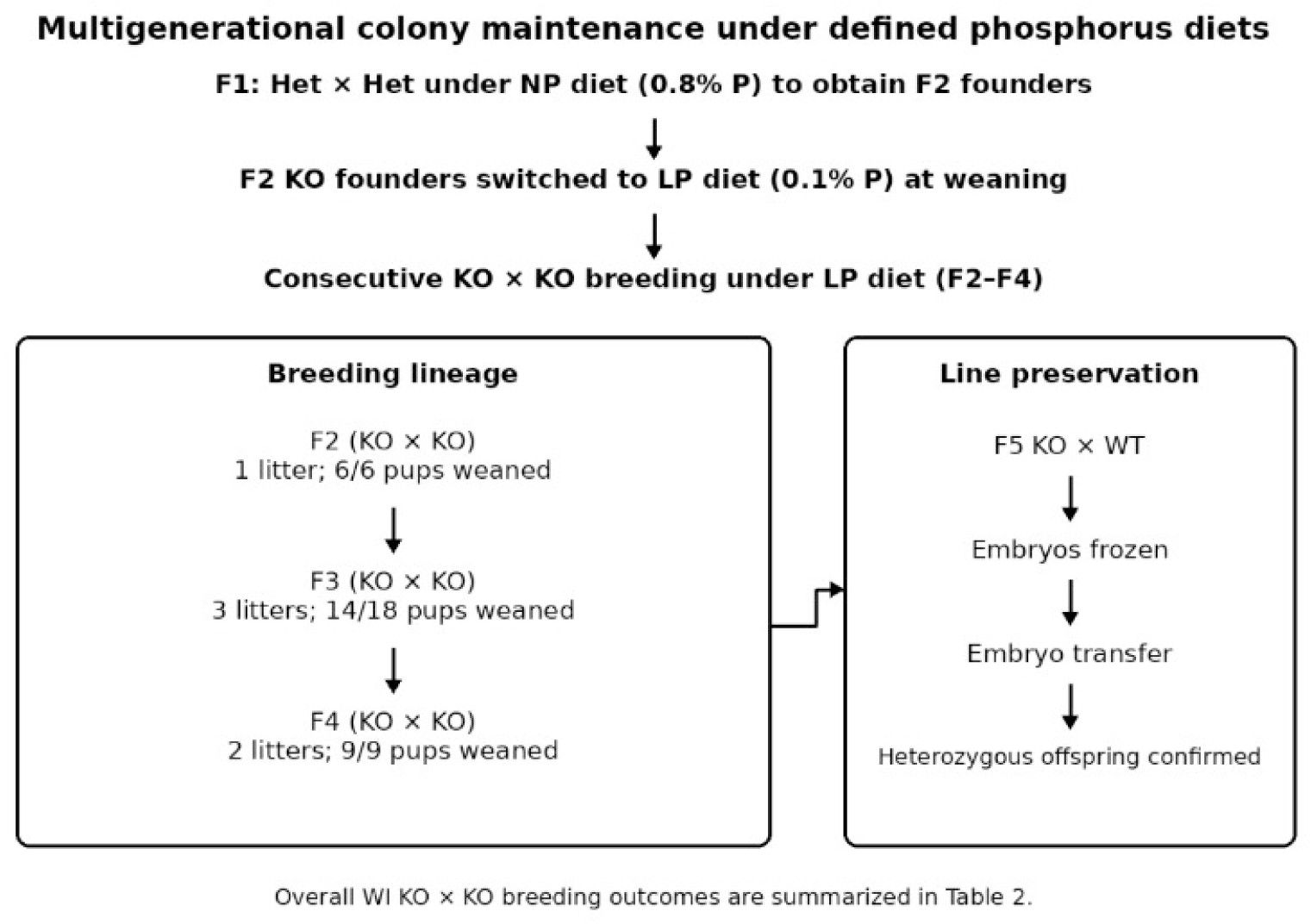
Multigenerational maintenance of WI KO colonies under LP conditions. Colony propagation showing natural KO × KO breeding under LP conditions across generations. Numbers indicate pups born and surviving to weaning within the illustrated lineage; aggregate breeding outcomes are summarized in Table 2. KO colonies were maintained to at least F5. NP denotes the normal-phosphorus diet (0.8% phosphorus), and LP denotes the low-phosphorus diet (0.1% phosphorus). Maternal LP feeding began at mating and continued through gestation and lactation; LP feeding was continued after weaning.

**Table 2.** Natural KO × KO breeding outcomes in LP-maintained WI rats.

| Strain | Litters | Born | Weaned | Survival (%) |
| --- | --- | --- | --- | --- |
| WI α-Klotho | 15 | 85 | 74 | 87.1 |

Vaginal smear analysis in LP-maintained KO females showed repeated estrous-type smears in five of six animals, with intervals broadly consistent with an approximately 4-day cycle (Fig. 9). Some intervals were 8 days. Because sampling was not performed on weekends and holidays, these longer intervals could reflect missed observations or altered cyclicity and could not be distinguished conclusively. In the remaining animal, only one estrous smear was observed, and cyclicity could not be clearly determined.

**Figure 9.**
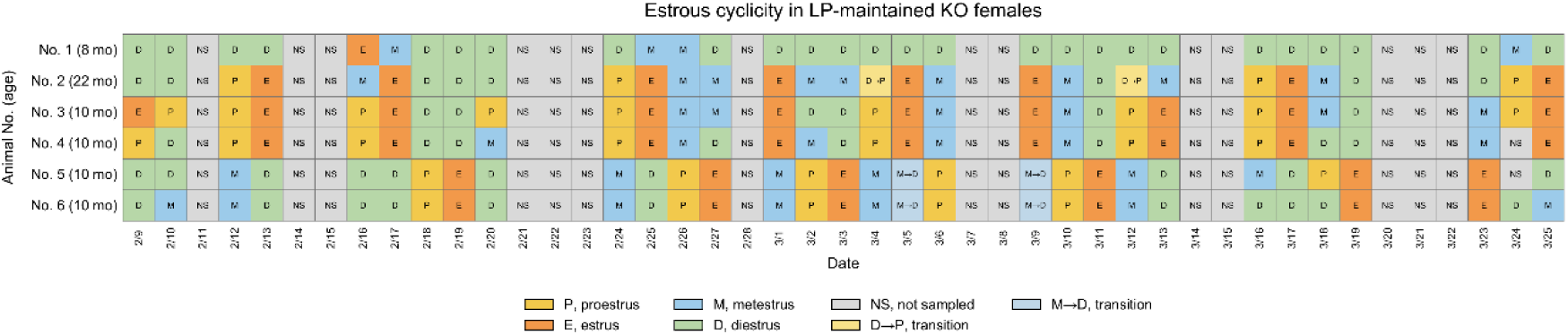
Estrous cyclicity in LP-maintained KO females. Vaginal smear analysis showing estrous cycle stages over time. Repeated estrous-type smears at approximately 4-day intervals were observed in most animals. Days without sampling, including weekends and holidays, were recorded as not sampled. Longer intervals could reflect missed observations or altered cyclicity and were not interpreted as definitive evidence for either explanation.

Timed mating after confirmation of proestrus resulted in successful delivery in several animals. Collectively, these data demonstrate that LP feeding restores sufficient gonadal maturation and reproductive capacity to enable natural KO × KO breeding.

### Heterogeneous 24-h metabolic rhythms in LP-maintained KO rats

We characterized metabolic profiles over the 24-h interval beginning at 19:00 on the recording start day (Fig. 10). WT animals under NP and LP conditions showed relatively high dark/light ratios for oxygen consumption (VO₂) and carbon dioxide production (VCO₂), consistent with nocturnal metabolic predominance. Het animals under NP conditions showed somewhat lower ratios, while dark-phase predominance was largely maintained under LP conditions.

**Figure 10.**
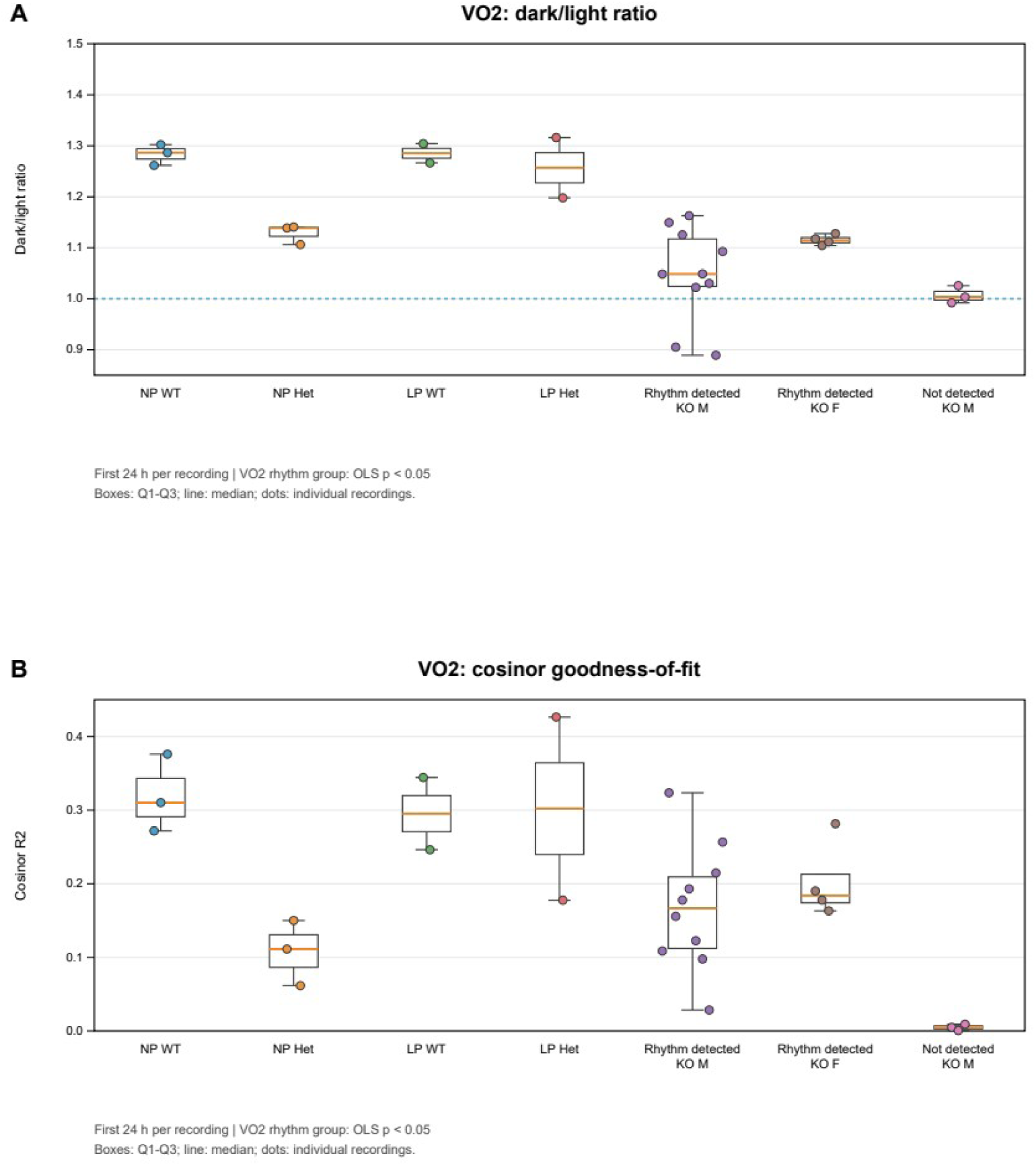
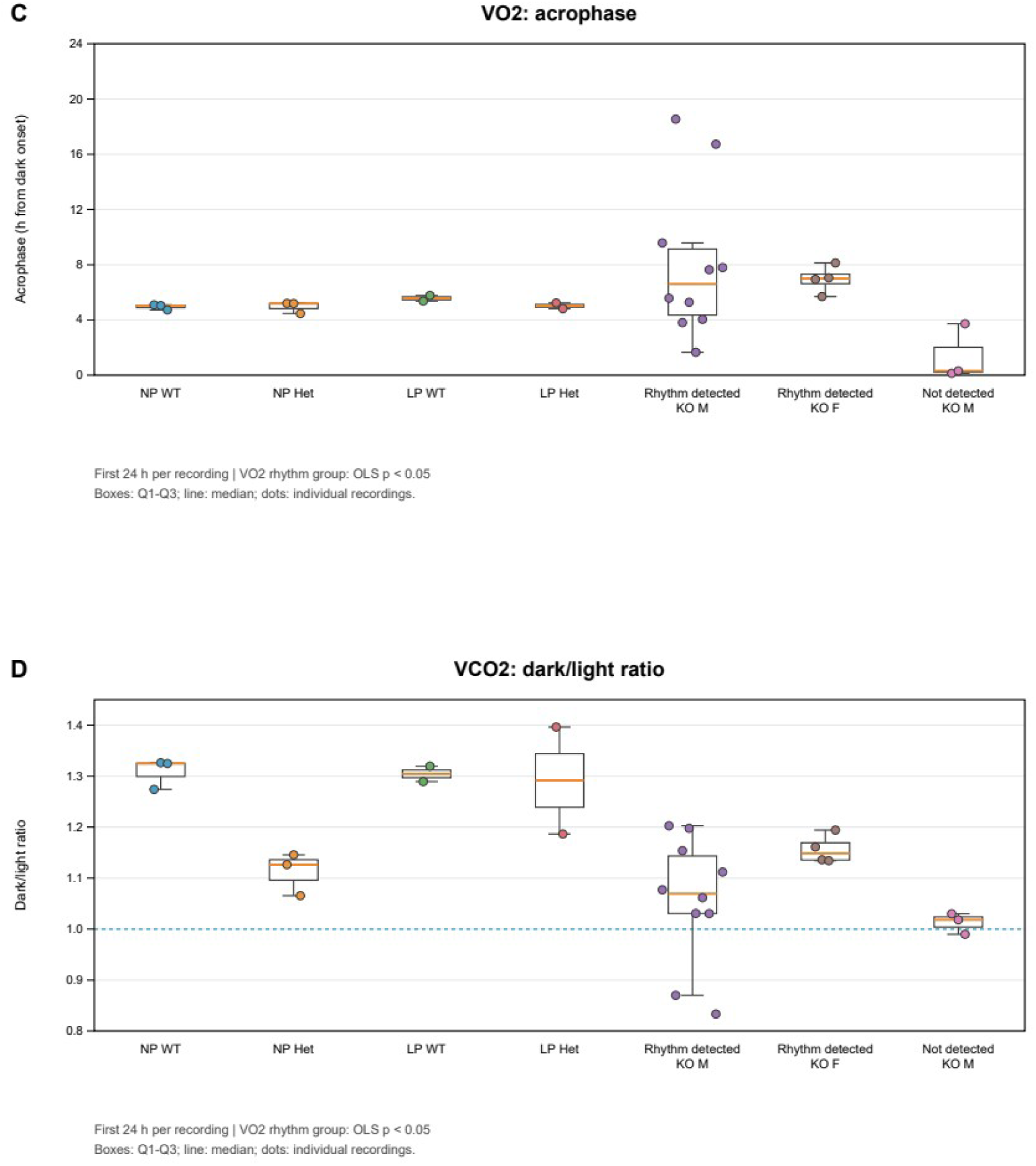
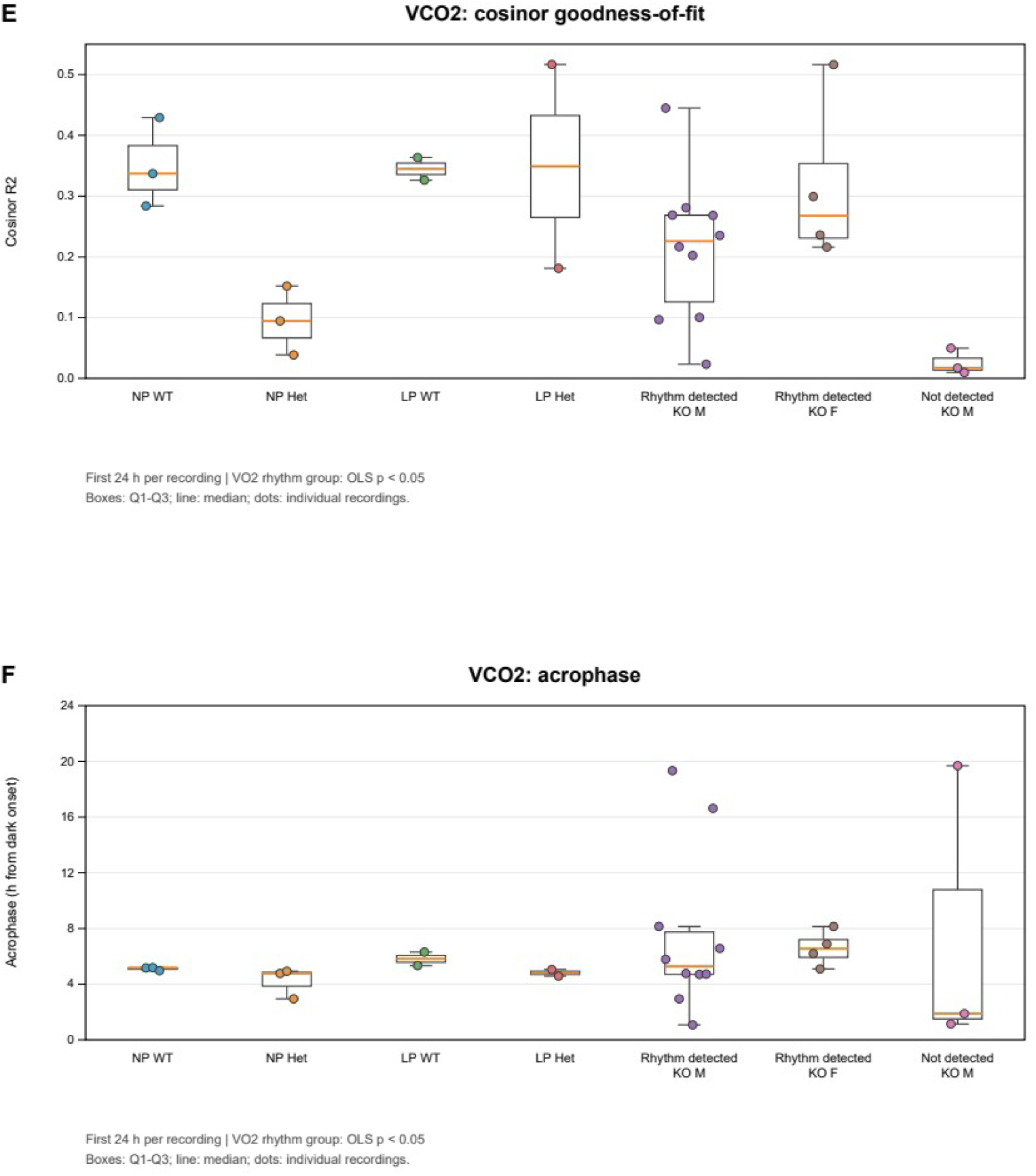
Heterogeneous 24-h metabolic profiles in LP-maintained KO rats. Profiles of oxygen consumption (VO₂) and carbon dioxide production (VCO₂), with dark/light ratios, cosinor goodness-of-fit (R²), and acrophase. WT and Het animals generally showed dark-phase predominance, whereas KO animals showed greater variability. Descriptions by VO₂ rhythmicity and sex are exploratory. KO animals with relatively clear VO₂ rhythms tended to retain dark-phase predominance, whereas individuals with weak or unclear rhythms showed attenuated oscillations, lower fit, and more variable phase estimates. These observations were obtained under a light–dark cycle.

LP-maintained KO animals showed heterogeneous profiles. In exploratory descriptions by sex and VO₂ rhythmicity, individuals with relatively clear rhythms showed VO₂ and VCO₂ dark/light ratios greater than 1, whereas males with weak or unclear rhythms had ratios approaching 1. Cosinor goodness-of-fit (R²) was generally lower in LP-maintained KO animals than in WT animals. In WT and Het animals, peak timing was distributed around approximately 5 h after dark onset. KO animals with relatively clear rhythms tended to have later phases, whereas phase estimates were more variable in males with weak or unclear rhythms.

Thus, while dietary LP restriction restores reproductive viability, 24-h metabolic rhythmicity remains heterogeneously altered in rescued KO animals. Exploratory mouse observations are provided in the Supplementary Results.

## Discussion

Dietary phosphorus restriction supported offspring recovery, reproductive development, and natural KO × KO breeding in WI rats carrying a large α-Klotho deletion, enabling maintenance to at least F5. The central advance is a practical dietary approach to maintaining a severe-deficiency rat model across generations. LP-maintained KO animals nevertheless retained heterogeneous 24-h metabolic profiles, showing that reproductive viability does not imply complete physiological recovery.

Maternal LP feeding was associated with higher overall survival to weaning, a higher proportion of KO animals among survivors, and reduced plasma phosphorus in KO pups. These observations are consistent with the importance of phosphate burden in α-Klotho-deficient states (Morishita et al., 2001; Ohnishi and Razzaque, 2010; Kuro-o, 2010; Razzaque, 2011). The LP group still deviated from the expected Mendelian ratio, and the reported goodness-of-fit tests do not directly test differences between dietary groups. Overall survival and KO recovery should therefore be distinguished from genotype-specific survival. Calcium increased under LP conditions, so the mineral changes cannot be interpreted as complete restoration of homeostasis. The contributions of gestation and lactation were not separated.

Continued LP feeding supported growth and gonadal development sufficiently for natural breeding. Histological and sperm observations provided additional evidence of reproductive development. WI KO × KO mating produced 85 pups across 15 litters, of which 74 survived to weaning, allowing multigenerational maintenance. Earlier studies demonstrated dietary improvement in hypomorphic Klotho-mutant mice and restoration of reproductive ability in Klotho/NaPi2a double-knockout mice (Morishita et al., 2001; Ohnishi and Razzaque, 2010). Our findings extend this framework to dietary maintenance of a large-deletion rat line. They do not establish that all aspects of growth or reproductive performance are normalized; pregnancy rates, mating success, and variation across breeding pairs require further characterization.

Despite successful breeding, metabolic profiles remained heterogeneous. Differences in body size, age, feeding behavior, and locomotor activity may contribute to the observed patterns, and feeding and activity were not directly measured. Sex and selection of animals for recording may also influence the profiles. Measurements under a light–dark cycle describe daily metabolic organization, and do not by themselves establish an intrinsic clock defect or a phosphate-independent mechanism. Stratification according to VO₂ rhythmicity should be interpreted descriptively, because the grouping criterion overlaps with some rhythm outcomes. Phase estimates in animals with weak rhythms also require caution. The ability to maintain these animals nevertheless creates an opportunity for controlled studies of residual metabolic phenotypes.

The deletion encompasses the proximal promoter region and exons 1–5, and deletion-junction sequencing provides a genetically defined model for long-term studies. The allele should be interpreted according to the available genomic evidence rather than assuming that every possible residual transcript or protein product has been excluded. The WI background provided a stable reproductive reference, whereas the limited observations in other rat backgrounds do not establish strain-independent rescue. The mouse observations involved different allele structures and were descriptive; they do not independently demonstrate a comparable dietary rescue effect.

Several limitations should be addressed in future work. Dietary exposure during gestation, lactation, and post-weaning was not separated into independent interventions. The survival summaries do not provide complete genotype-specific mortality, and litter-level dependence should be considered in comparisons of offspring outcomes. Sampling over P20–32 and the timing of weaning should be accounted for when interpreting mineral measurements. The dietary formulations differed, and effects of components other than phosphorus cannot be excluded without a complete composition comparison. The metabolic observations also require larger, appropriately matched groups and repeated measurements to assess reproducibility. These limitations define the scope of the present evidence without diminishing the demonstrated feasibility of natural KO × KO breeding and multigenerational maintenance.

Dietary phosphorus restriction from the maternal period through post-weaning enabled natural breeding and multigenerational maintenance of severe α-Klotho-deficient WI rats. Heterogeneous metabolic rhythms persisted, showing that successful colony maintenance does not imply complete physiological recovery. This maintainable model may support future genetic complementation studies.

## Materials and Methods

### Animals and ethics statement

Rats were maintained under standard specific pathogen-free conditions with free access to food and water. The light–dark cycle and room temperature were controlled according to institutional facility standards. All animal procedures were approved by the Institutional Animal Care and Use Committee of Jichi Medical

University (approval nos. 24038-03, 24040-02, 24123-03, 24124-02, 25005-01, 25006-01, 25007-01) and were conducted in accordance with the institutional guidelines for animal experimentation.

### Generation of α-Klotho-deficient rats

To generate α-Klotho-deficient rats, a large deletion spanning exons 1–5 of the rat α-Klotho gene was introduced using CRISPR/Cas9. For genome editing, ribonucleoprotein (RNP) complexes were prepared using guide RNAs from the Alt-R CRISPR-Cas9 System (Integrated DNA Technologies, IDT) together with TrueCut Cas9 Protein v2 (Thermo Fisher Scientific). Guide RNA sequences used were rKla_up_gRNA 5’-GCCAAACGTCGCGCAAAGTT-3’ and rKla_dw_gRNA 5’-GATCGCTGTCGGGTCTACTA-3’. The RNP complexes were introduced into one-cell or two-cell embryos by electroporation using a Nepa Gene electroporator. Embryos were handled in PB1 medium and cultured in KSOM Rat medium supplemented with Y-27632. Embryos that had developed were transferred into the oviducts of pseudopregnant recipient females. Mutant lines were established on Wistar-Imamichi (WI), F344, and Wistar-Kyoto (WKY) backgrounds. The use of Y-27632 during embryo handling followed the temporary-treatment-medium approach described by Nagao et al. (WO2021193596A1; Japanese Patent No. 7722712 [JP7722712B2]).

### Genotyping

Genotyping was performed using a combination of three PCR assays targeting the 5’ wild-type allele (yielding an ∼800 bp product), the 3’ wild-type allele (∼600 bp), and the deletion junction (∼800 bp). This strategy allowed clear discrimination among wild-type (WT), heterozygous (Het), and knockout (KO) animals. Formation of the intended deletion junction was further confirmed by Sanger sequencing of the deletion-junction PCR product. The primers used were KLAupF1 primer 5’-CAGGGAAAGGAGAAACAGAACC-3’, KLAupF3 primer 5’-GGGGTCTAATTAAGGAACGGATT-3’, KLAupR1 primer 5’- AGCAGATGCAGCGACAGCAAAC-3’, KLAdwF2 primer 5’-AGTGATTTGGTTGAACGTCCTTT-3’, KLAdwR2 primer 5’-CATTCATGAGGAAAACCATACCA-3’. The deletion design included the proximal promoter region so that the knockout allele could be identified directly by deletion-junction PCR.

### Dietary conditions

Animals were maintained on a normal-phosphorus diet (NP; MFG, Oriental Yeast Co., Ltd., Tokyo, Japan; 0.8% phosphorus) or a low-phosphorus diet consisting of a modified TD.10662 formulation (LP; Oriental Yeast Co., Ltd.; 0.1% phosphorus). Maternal phosphorus restriction was defined as LP feeding from the start of mating through gestation and lactation. Post-weaning phosphorus restriction was defined as continuation of LP feeding after weaning. NP animals were maintained on MFG during the corresponding experimental period.

### Assessment of survival and genotype distribution at weaning

WI heterozygous intercrosses were performed under NP or LP conditions. For the LP group, females were switched to LP feeding at the start of mating and maintained on LP throughout gestation and lactation. Litter size, the number born, survival to weaning, and the genotype distribution among survivors were recorded. Overall survival to weaning was calculated as the number weaned divided by the number born, multiplied by 100. Genotype percentages used the total number of weaned survivors as the denominator. Deviation from the expected Mendelian 1:2:1 ratio was assessed by chi-square goodness-of-fit testing.

### Blood biochemistry

Blood samples were collected at postnatal days 20–32. Plasma phosphorus, calcium, and magnesium concentrations were measured using a dry chemistry system. NP and LP groups were compared within each genotype using two-sided Mann–Whitney U tests, with Holm correction for multiple comparisons.

### Tissue collection

Animals were anesthetized with isoflurane and euthanized after loss of withdrawal reflexes in response to limb and tail pinch had been confirmed. Kidneys were collected after removal of the renal capsule. Testes were dissected free of the caput and cauda epididymides and surrounding adipose tissue. Ovaries were collected after removal of the ovarian bursa. Tissues were weighed and fixed in Bouin’s solution or 10% formalin.

### Histology

Fixed tissues were dehydrated, cleared in xylene, and embedded in paraffin. Sections of 4–5 µm thickness were prepared using a rotary microtome (Leica) and stained with hematoxylin and eosin (H&E). Sections were examined and imaged using an upright microscope (Olympus).

### Scanning electron microscopy

Collected sperm were incubated in human tubal fluid (HTF) medium and fixed in 0.1 M cacodylate buffer containing 3.5% glutaraldehyde. After drying, samples were coated with platinum-palladium and examined using a scanning electron microscope (Hitachi).

### Evaluation of estrous cyclicity

Estrous cyclicity in LP-maintained KO females was evaluated by vaginal cytology. Smears were classified into proestrus, estrus, metestrus, and diestrus. Days on which sampling was not possible, including weekends and holidays, were recorded as “not sampled.” When appropriate, timed mating was performed after confirmation of proestrus to assess the establishment of pregnancy.

### Indirect calorimetry

Whole-body respiratory metabolism was measured using a small-animal metabolic monitoring system (MK-5100MSII; Muromachi Kikai, Tokyo, Japan). Each rat was housed individually in a rat chamber (300 × 190 × 250 mm) and acclimated to the chamber environment before measurement. Measurements were performed at 25 °C under a 12:12-h light-dark cycle (light, 07:00–19:00; dark, 19:00–07:00). Oxygen consumption (VO₂), carbon dioxide production (VCO₂), and respiratory quotient (RQ) were recorded continuously. RQ was calculated as VCO₂/VO₂. Energy expenditure was calculated as E = (1.07 × RQ + 3.98) × VO₂. Depending on the analysis, these parameters were evaluated as time-averaged values, light- and dark-phase averages, or circadian rhythm parameters.

### Cosinor analysis

The 24-h rhythmicity of metabolic variables under the light–dark cycle was assessed using single-component cosinor analysis with a fixed 24-h period, following established rhythmometry methods (Cornelissen, 2014). The mesor, amplitude, and acrophase were estimated by ordinary least-squares regression. Differences between light- and dark-phase means were also tested. Individuals showing significant light– dark differences and significant 24-h rhythmicity were classified as rhythm-positive. Descriptions stratified by VO₂ rhythmicity and sex were treated as exploratory.

Analyses were performed in Python 3.13.2 using NumPy 2.4.0, pandas 2.3.3, SciPy 1.17.0, statsmodels 0.14.6, and matplotlib 3.10.8.

### Statistical analysis

Deviation of genotype frequencies at weaning from the expected Mendelian ratio of 1:2:1 was assessed using the chi-square goodness-of-fit test. Plasma mineral concentrations were compared between NP and LP groups within each genotype using two-sided Mann–Whitney U tests with Holm correction for multiple comparisons. Reported genotype goodness-of-fit tests were not treated as direct comparisons between dietary groups.

Body weight at postnatal day 21 was analyzed using two-way analysis of variance, with genotype and diet as fixed factors and including the genotype × diet interaction. Pairwise group comparisons were performed using Tukey’s multiple-comparisons test.

The 24-h rhythmicity of metabolic variables was assessed using single-component cosinor analysis with a fixed 24-h period based on ordinary least-squares regression. Group comparisons of metabolic variables and cosinor-derived parameters were performed using Welch’s t-test or the Mann–Whitney U test, as specified for the corresponding analyses.

For box-and-whisker plots, the horizontal line within each box represents the median, and the lower and upper boundaries of the box represent the first and third quartiles, respectively. The whiskers extend to the most extreme observations within 1.5 times the interquartile range below the first quartile and above the third quartile. All individual observations, including values beyond the whiskers, are shown as data points. Mean values are not shown unless otherwise stated. The value of n represents the number of animals analyzed.

Unless otherwise stated, tests were two-sided and P < 0.05 was considered statistically significant.

## Acknowledgements

We sincerely thank the members and staff of the Center for Experimental Medicine, Jichi Medical University, the staff of the Institute of Immunology Co., Ltd., and the members of the Division of Functional Biochemistry, Department of Biochemistry, Jichi Medical University, for their valuable support and assistance in this study. We also thank Dr. Ken-ichiro Ishida of the Faculty of Life and Environmental Sciences, University of Tsukuba, for his guidance on scanning electron microscopy.

## Data and Resource Availability

Data underlying the findings reported in this study are available from the corresponding author upon reasonable request, subject to applicable institutional requirements. Availability of the α-Klotho-deficient animal resources for collaborative studies may be discussed with the corresponding author and is subject to institutional approval and applicable material-transfer requirements.

## Collaboration Statement

This preprint reports the current stage of an ongoing research program. Further mechanistic and physiological characterization of these α-Klotho-deficient models is ongoing. We welcome inquiries from investigators interested in collaborative studies using these models, including studies of mineral metabolism, renal and vascular pathology, reproductive physiology, circadian metabolism, and genetic or therapeutic rescue. Collaborative use of the animal resources will be considered subject to applicable institutional approvals and material-transfer requirements.

## Supplementary Results

### Additional rat backgrounds and exploratory growth observations

Deletion-junction sequences were confirmed in two F344 lines and one WKY line. Breeding observations used one line in each background. Under NP conditions, F344 and WKY heterozygous intercrosses yielded 82/88 and 45/53 weaned survivors, respectively. Their genotype distributions did not significantly deviate from 1:2:1 (Supplementary Table S1). LP observations comprised one litter per background and cannot establish reproducible dietary rescue or differences between backgrounds. A single F344 KO × KO litter under LP conditions yielded four pups, all of which survived to weaning (Supplementary Table S2).

Body weight observations in F344 and WKY are presented descriptively in Supplementary Fig. S1. Ages differed between groups, and some groups were small. Supplementary Fig. S2 presents KO body weights normalized to mean WT body weight within each background and dietary condition. The WKY LP group lacked WT animals, preventing calculation of a corresponding KO/WT ratio. These observations are not used to establish an independent post-weaning effect or a strain-independent rescue response.

### Exploratory mouse observations

A C57BL/6J (B6) mouse line carried a large deletion spanning exons 1–5. In Mus spretus-related editing, an exon 2–3′UTR deletion had been intended, but the recovered line instead carried an exon 2-restricted 4-bp mutation predicted to cause a frameshift and premature truncation. This line is described as a frameshift mutant rather than a complete genomic-deletion KO. Editing designs and sequence confirmation are shown in Supplementary Figs. S3 and S4. Six B6 and three Mus spretus-related lines were sequence-confirmed; two B6 and one Mus spretus-related line were used for breeding observations.

Under LP-supported conditions, mutant animals reached weaning in the mouse lines examined. B6 KO females showed estrus, pregnancy, and delivery, and B6 KO males showed evidence of fertilizing ability. B6 KO animals maintained on NP feeding from the maternal period were not maintained beyond 12 weeks. Spermatogenesis was observed in homozygous Mus spretus-related frameshift-mutant males. These observations lacked detailed quantitative breeding comparisons and do not independently establish the effect of LP feeding. Exploratory metabolic profiles in B6 mice are shown descriptively in Supplementary Fig. S5.

## Supplementary Methods

### Supporting rat and mouse line generation

In addition to WI, α-Klotho deletion lines were established on F344 and WKY rat backgrounds using the rat editing strategy described above. During embryo production without Y-27632, either no pups were obtained or recovered pups lacked the KO-specific deletion band. When Y-27632 was included in the culture medium, animals carrying the intended deletion band were recovered. These are procedural observations rather than a quantified test of inhibitor efficacy.

### Generation and genotyping of supporting mouse α-Klotho mutant lines B6 exon 1–5 deletion line

To generate a severe α-Klotho-deficient mouse line on the C57BL/6J background, a large deletion spanning exons 1–5 of the mouse α-Klotho gene was introduced by CRISPR/Cas9-mediated genome editing. Two guide RNAs were designed to target the 5′ and 3′ flanking regions of the deletion interval. The guide RNA sequences were Mkla5gRNA, 5′-GAAACTTCCAATCGGGCAGA-3′, and Mkla3gRNA, 5′-CGCAGTTTCAGTGCATCGAC-3′. According to the design, the intended deletion size was approximately 66 kb. Genotyping was performed using three PCR assays: a 5′ wild-type allele PCR, a 3′ wild-type allele PCR, and a deletion-junction PCR specific for the edited allele. For the 5′ wild-type allele, primers Mkla5F (5′-GGTAGAGGCGATGGAGGAG-3′) and Mkla5R (5′-CCCTTCACACTCTATCCAGTC-3′) were used to amplify an approximately 800-bp product. For the 3′ wild-type allele, primers Mkla3F (5′-GCTTGGAGGTGCTAGCTAG-3′) and Mkla3R (5′-CCAGGTGATGAAGCCTGGTC-3′) were used to amplify an approximately 700-bp product. For the deletion-junction allele, primers Mkla5F and Mkla3R were used to amplify an approximately 600-bp product spanning the edited junction. Formation of the intended deletion junction was confirmed by sequence analysis of the deletion-junction PCR product.

### Mus spretus-related exon 2 mutant line

Mus spretus was included as a physiologically distinct background because of its relatively low water intake and its relevance for comparison of renal function with B6 mice. These considerations motivated exploratory observations rather than a controlled background comparison.

For the Mus spretus-related α-Klotho editing experiments, the Mus spretus line used for genome editing was maintained on a Tyr-/-albino background. In addition, BSF1 (C57BL/6J × Mus spretus) animals used as recipients were also Tyr-/- and albino.

To generate a Mus spretus-related α-Klotho mutant line, CRISPR/Cas9-mediated editing was designed to delete the region spanning exon 2 to the 3′ untranslated region (3′UTR) of the α-Klotho gene. Two guide RNAs were designed to target exon 2 and the 3′UTR, respectively. The guide RNA sequences were Ex2gRNA, 5′-TAACATGAAGTTCCGCCAAT-3′, and 3UTRgRNA, 5′-GGTAGATCTCGGCGGCCGCA-3′.

According to the design, the intended deletion size was approximately 12 kb. For this line, genotyping was designed using three PCR assays: an exon 2 wild-type allele PCR, a 3′UTR wild-type allele PCR, and a deletion-junction PCR. For the exon 2 wild-type allele, primers Mklaex2_5F (5′-TCCTCGAAGAATGACCGACT-3′) and Mklaex2_5R (5′-TGGTTCCCGAGACAAACCAG-3′) were used to amplify an approximately 300-bp product. For the 3′UTR wild-type allele, primers Mkla_3UTRF (5′-TTCTTAGTCGATCCGGCAGC-3′) and Mkla_3UTRR (5′-CTGTGCTACACCTGAGTGGT-3′)were used to amplify an approximately 300-bp product. For the deletion-junction assay, primers Mklaex2_5F and Mkla_3UTRR were designed to amplify an approximately 500-bp product spanning the edited junction.

However, in the recovered line used for analysis, a large deletion spanning exon 2 to the 3′UTR was not obtained. Instead, sequence analysis of the exon 2 amplicon identified a mutation confined to exon 2. Comparison with the reference Mus spretus sequence indicated a 4-bp alteration within exon 2, which was predicted to cause a frameshift and premature truncation of the encoded protein. Thus, although the original design targeted a broader exon 2–3′UTR interval, the line used in this study carried an exon 2-restricted frameshifting mutation.

## Supplementary Tables

**Supplementary Table S1.** Descriptive breeding outcomes in F344 and WKY heterozygous intercrosses.

| Strain | Diet | Litters | Born | Weaned | Overall survival to weaning (%) | Died | WT | Het | KO | $\chi^2$ | P value |
| --- | --- | --- | --- | --- | --- | --- | --- | --- | --- | --- | --- |
| F344 α-Klotho | NP | 11 | 88 | 82 | 93.2 | 6 | 19 | 49 | 14 | 3.73 | 0.155 |
| F344 α-Klotho | LP | 1 | 7 | 7 | 100.0 | 0 | 1 | 4 | 2 | 0.43 | 0.807 |
| WKY α-Klotho | NP | 6 | 53 | 45 | 84.9 | 8 | 13 | 18 | 14 | 1.84 | 0.398 |
| WKY α-Klotho | LP | 1 | 8 | 8 | 100.0 | 0 | 0 | 4 | 4 | 4.00 | 0.135 |

Counts of litters, pups born, and survivors under NP and LP conditions. Overall survival was calculated as Weaned / Born × 100. WT, Het, and KO counts represent all weaned survivors genotyped by PCR. Chi-square goodness-of-fit tests assess deviation from 1:2:1 within each dietary group and do not directly compare groups. LP observations were limited to one litter in each background; the data are
descriptive and do not establish reproducible rescue. Mortality before genotyping limits interpretation of genotype-specific survival.

**Supplementary Table S2.** Single-litter F344 KO × KO breeding observation under LP conditions.

| Strain | Litters | Born | Weaned | Survival (%) |
| --- | --- | --- | --- | --- |
| F344 α-Klotho | 1 | 4 | 4 | 100.0 |

One F344 KO × KO litter produced four pups, all of which survived to weaning. The observation is descriptive and does not establish reproducibility in this background.

## Supplementary Figures and legends

**Supplementary Figure S1.**
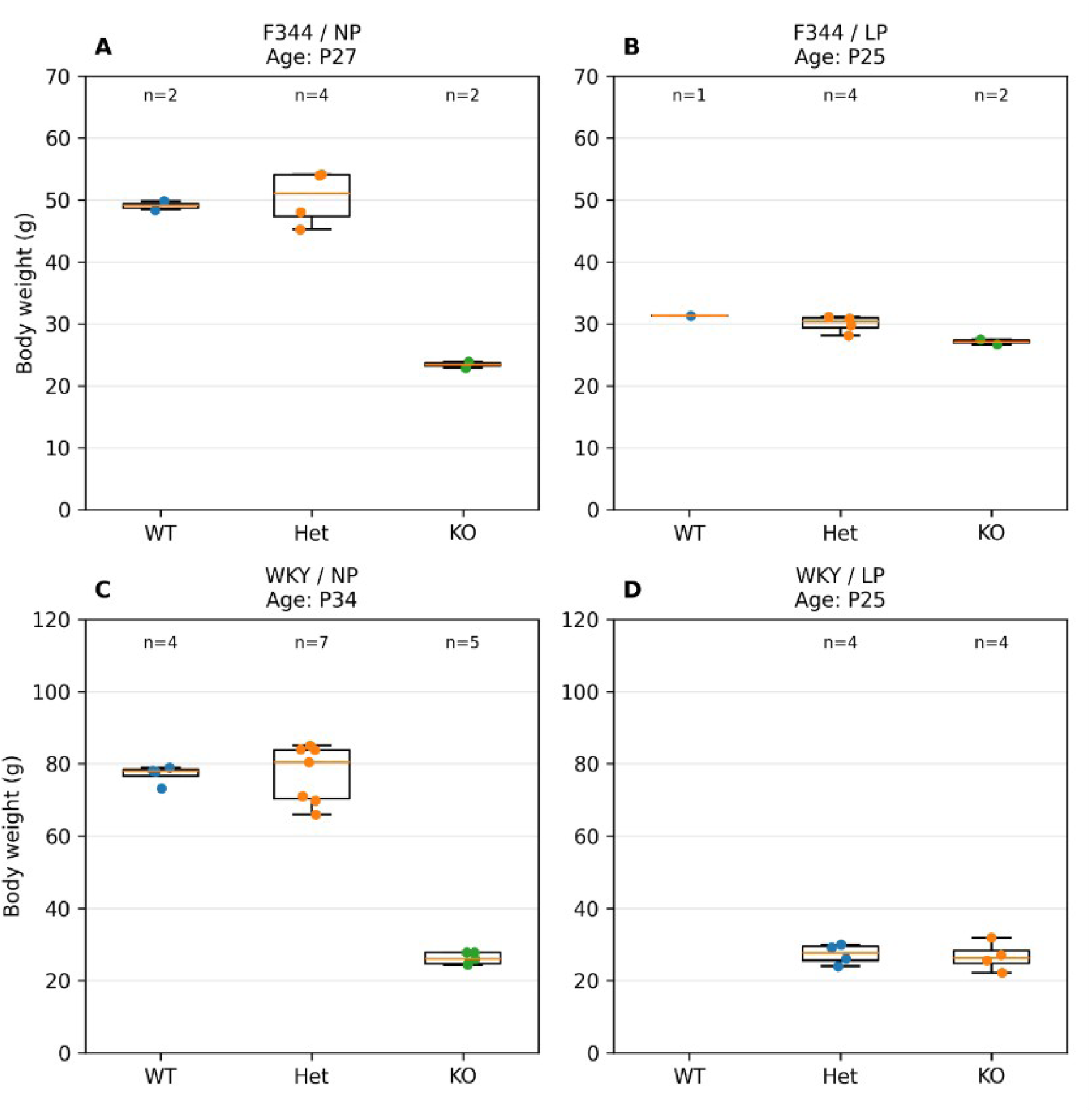
Descriptive body weights in F344 and WKY rats under NP and LP conditions. Body weights of WT, Het, and KO animals in F344 and WKY backgrounds. Each point represents one animal. Boxes show the median and interquartile range; whiskers extend to the most extreme observations within 1.5 × IQR. Individual observations and group sizes are shown. Sampling ages are indicated in each panel. A and B show F344 under NP and LP conditions, respectively; C and D show WKY under the corresponding conditions. Ages differed between groups and some groups were small. These data are descriptive and are not used to establish a strain-independent intervention effect. Y-axis scales were adjusted within each background.

**Supplementary Figure S2.**
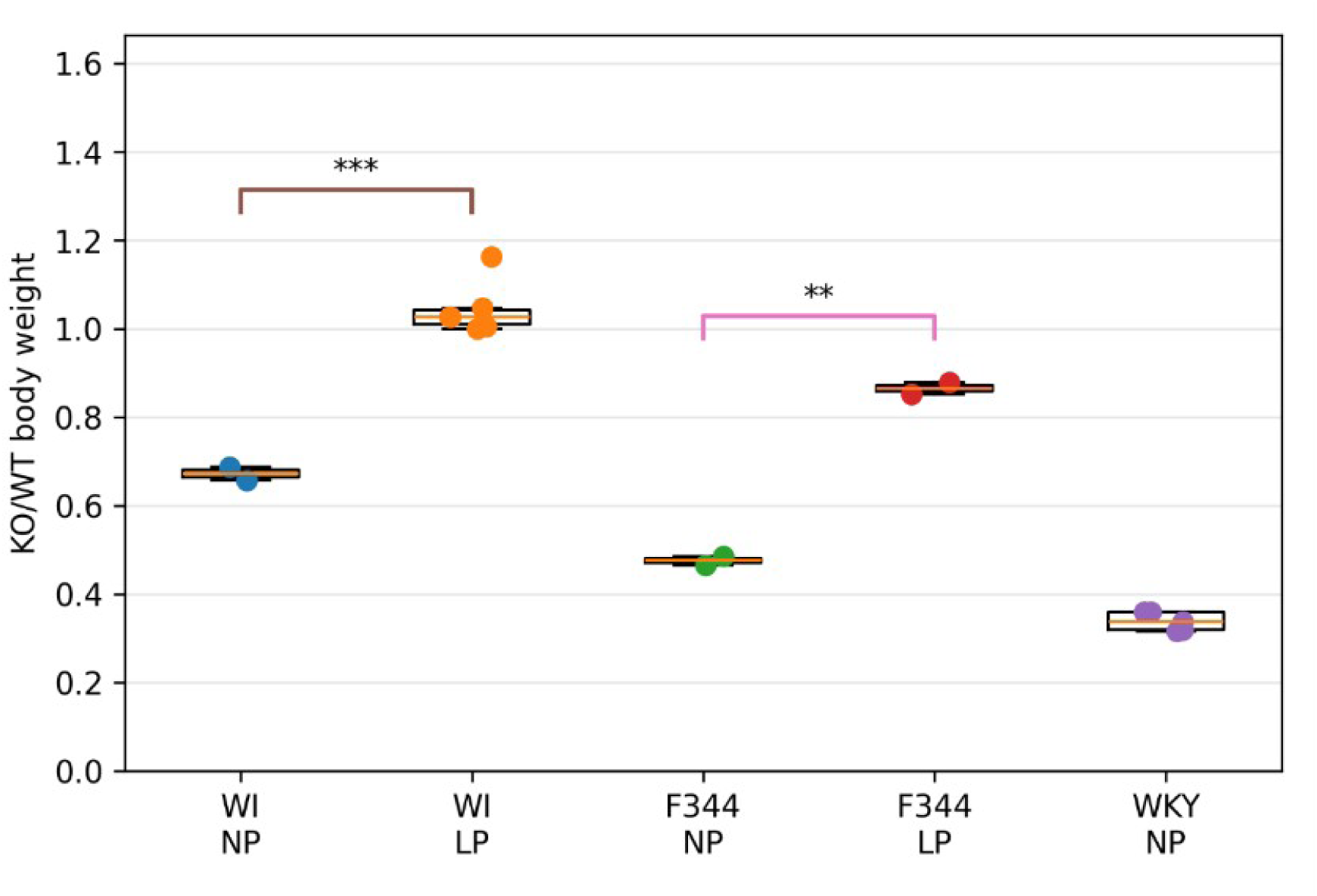
Descriptive KO body weight normalized to WT. KO body weight was normalized to mean WT body weight within each background and dietary condition. Each point represents one KO animal. Boxes show the median and interquartile range; whiskers extend to the most extreme observations within 1.5 × IQR. The original exploratory NP–LP comparisons used unpaired t-tests within each background. *P < 0.05, **P < 0.01, ***P < 0.001; ns, not significant. No KO/WT ratio was available for WKY LP because WT animals were absent, so WKY NP alone is not evidence for an NP–LP comparison. Age differences and small groups limit interpretation

**Supplementary Figure S3.**
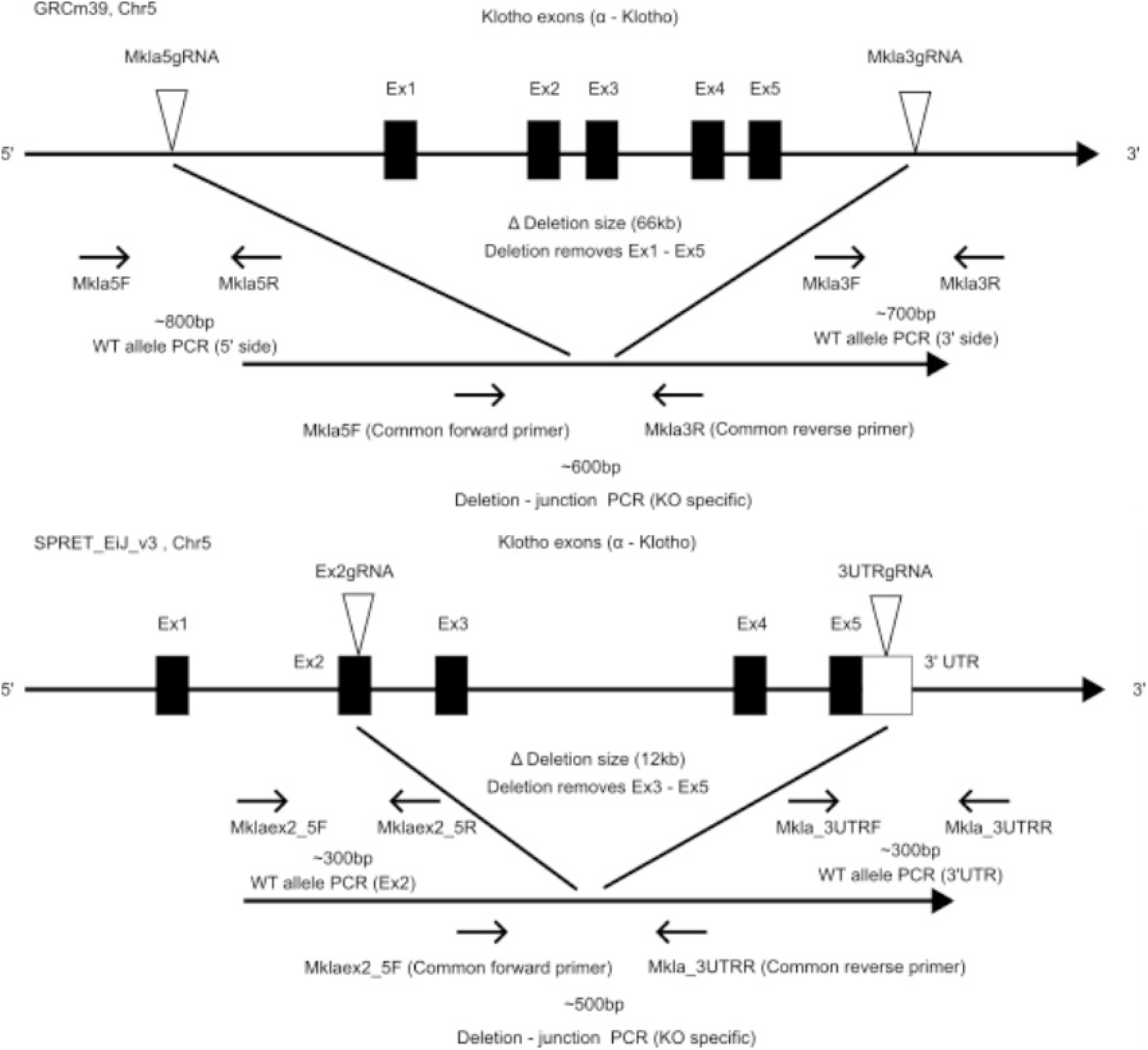
Genome-editing designs of exploratory mouse α-Klotho mutant lines. a, Design for deletion of exons 1–5 in B6 mice, with guide RNA target sites and wild-type allele and deletion-junction PCR strategies. b, Mus spretus-related editing design targeting exon 2 to the 3′UTR. This is the intended design: the recovered line carried an exon 2-restricted 4-bp frameshift mutation rather than the intended large deletion. Sequence confirmation is provided in Supplementary Fig. S4.

**Supplementary Figure S4.**
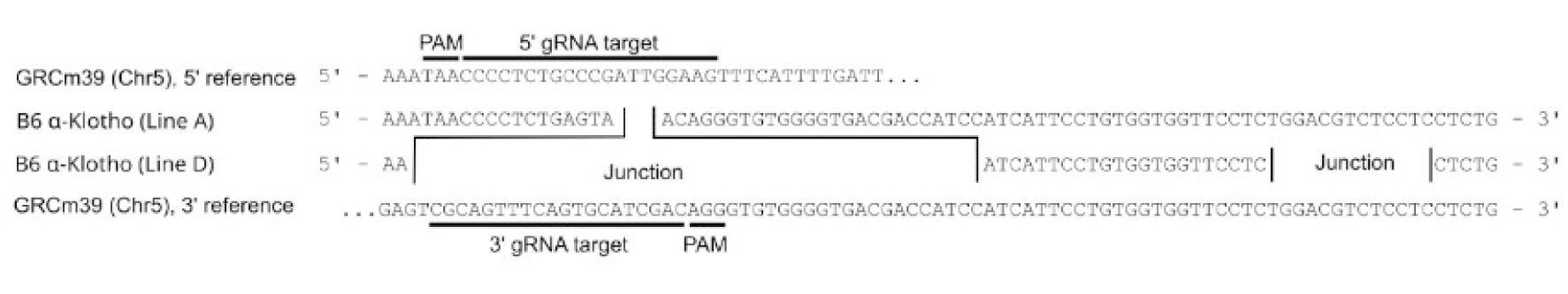
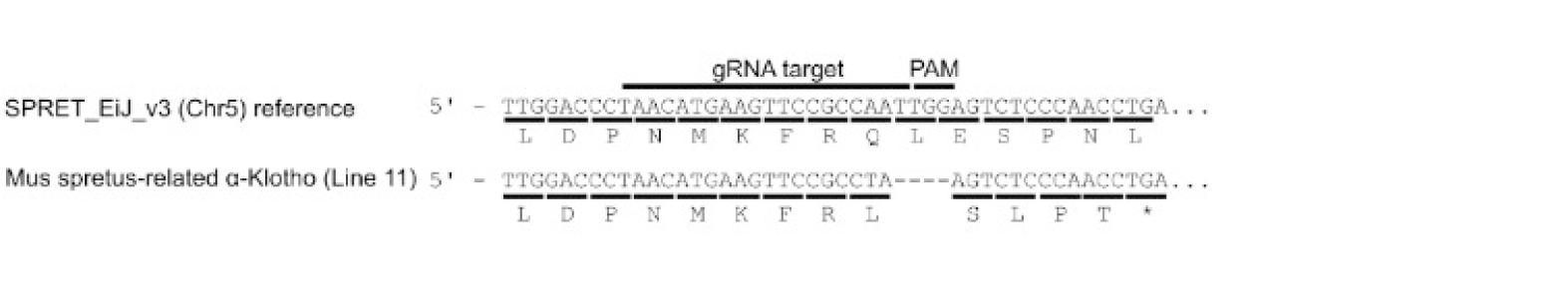
Sequence confirmation of exploratory mouse α-Klotho mutant alleles. a, Deletion-junction sequences in B6 exon 1–5 deletion lines. Six B6 lines were sequence-confirmed and two were used for breeding observations. b, Sequence confirmation of the Mus spretus-related allele. The recovered line carried an exon 2-restricted 4-bp mutation predicted to cause a frameshift and premature truncation, rather than the intended exon 2–3′UTR deletion. Three Mus spretus-related lines were sequence-confirmed and one was used for breeding observations.

**Supplementary Figure S5.**
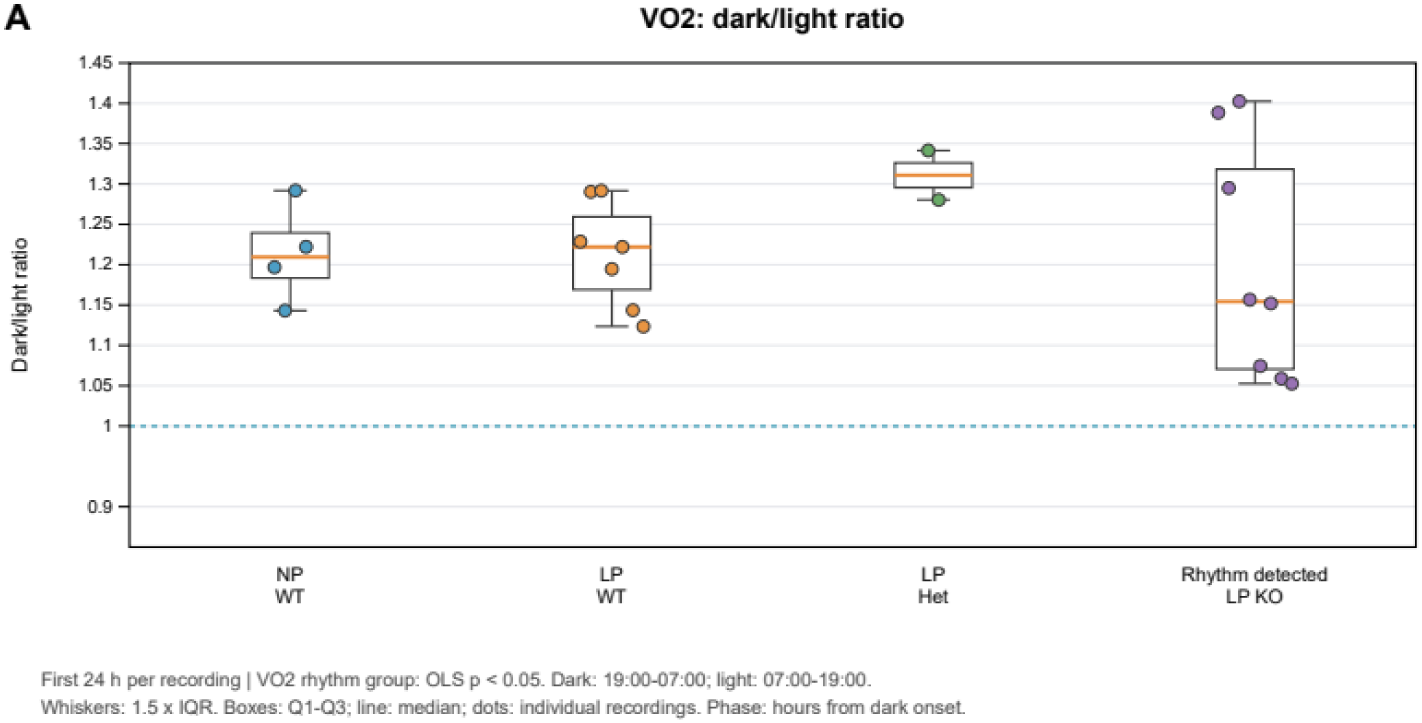
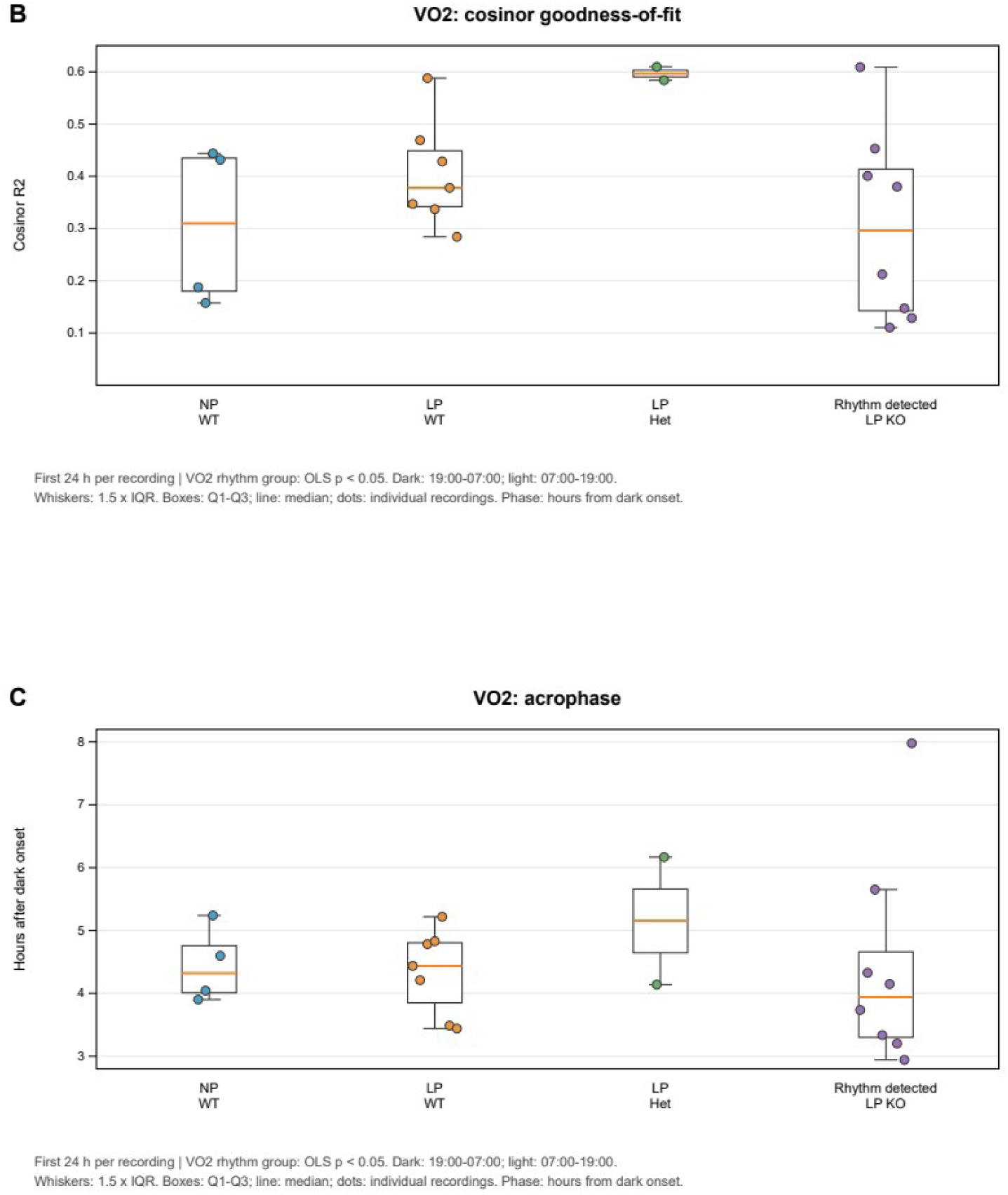
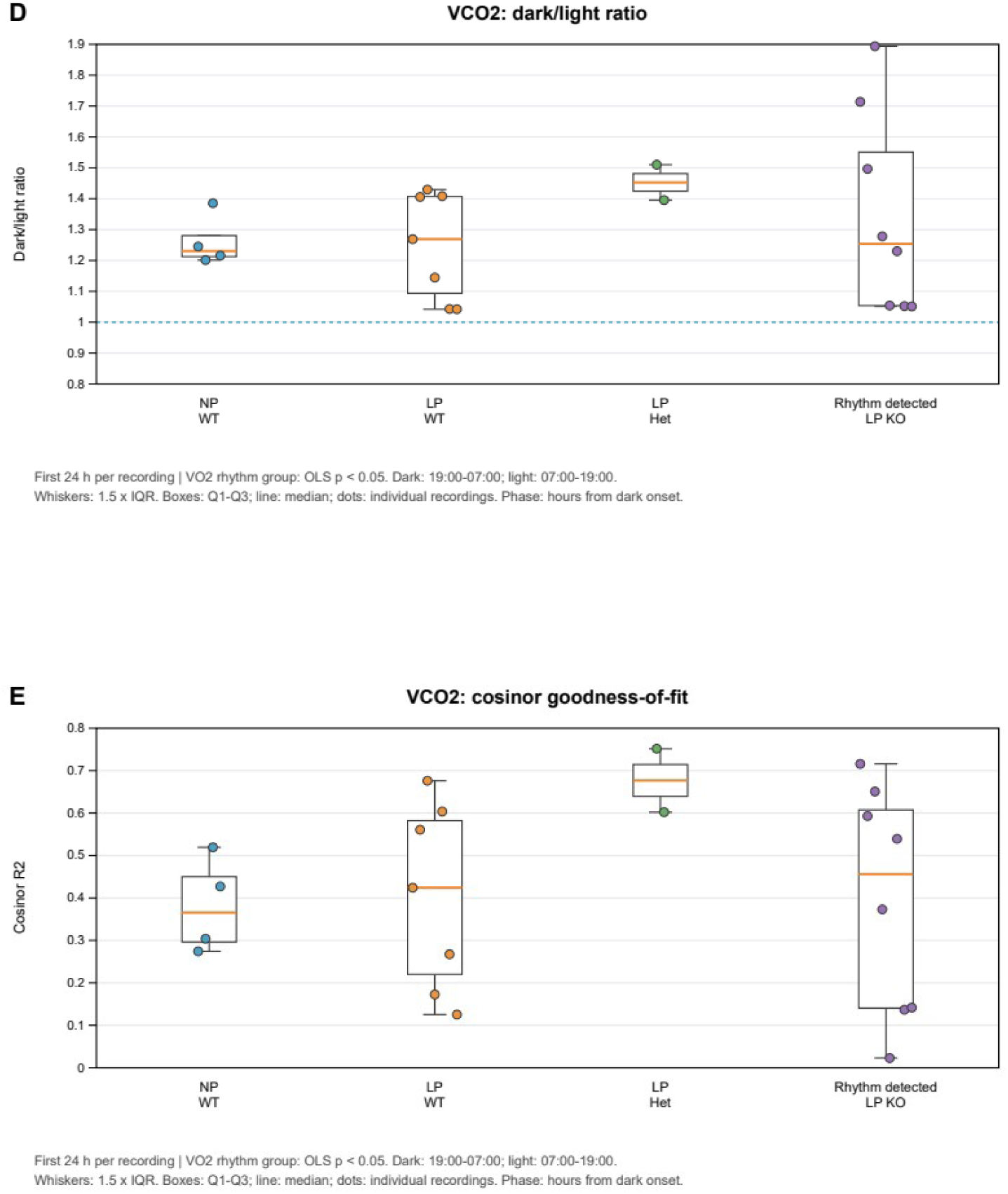
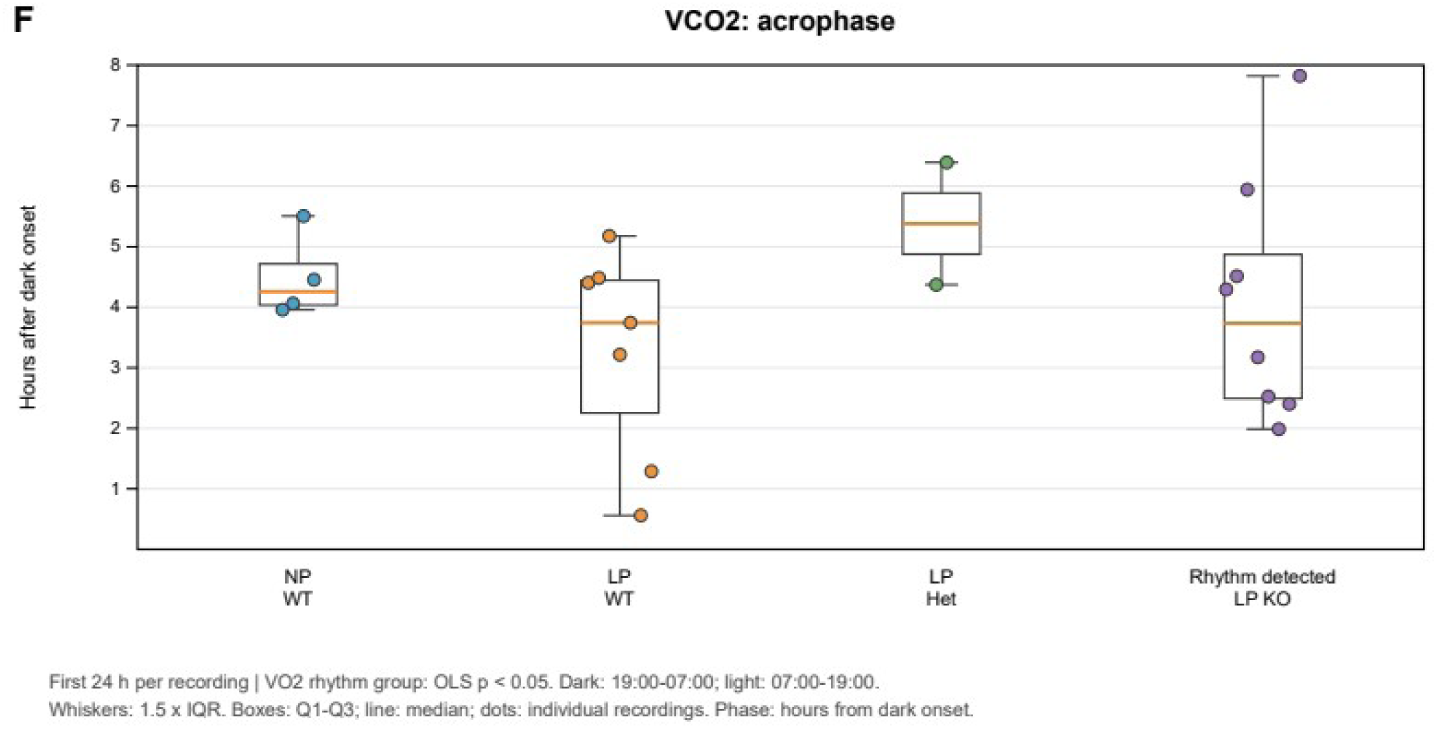
Descriptive 24-h metabolic indices in B6 α-Klotho mutant mice. Upper panels show VO₂-based indices and lower panels show VCO₂-based indices. Dark/light ratio, cosinor goodness-of-fit (R²), and acrophase are presented for exploratory description. Sample sizes were limited, and these data were not used for formal quantitative comparison in the main text.

## References

1. Shimada, T., et al. (2004). Targeted ablation of Fgf23 demonstrates an essential physiological role of FGF23 in phosphate and vitamin D metabolism. Journal of Clinical Investigation, 113, 561–568.

2. Kurosu, H., et al. (2006). Regulation of fibroblast growth factor-23 signaling by klotho. Journal of Biological Chemistry, 281(10), 6120–6123.

3. Urakawa, I., et al. (2006). Klotho converts canonical FGF receptor into a specific receptor for FGF23. Nature, 444, 770–774.

4. Nakatani, T., et al. (2009). In vivo genetic evidence for klotho-dependent, fibroblast growth factor 23 (Fgf23)-mediated regulation of systemic phosphate homeostasis. FASEB Journal, 23, 433–441.

5. Hu, M. C., et al. (2010). Klotho: a novel phosphaturic substance acting as an autocrine enzyme in the renal proximal tubule. FASEB Journal, 24, 3438–3450.

6. Kuro-o, M., et al. (1997). Mutation of the mouse klotho gene leads to a syndrome resembling ageing. Nature, 390, 45–51.

7. Ohnishi, M., and Razzaque, M. S. (2010). Dietary and genetic evidence for phosphate toxicity accelerating mammalian aging. FASEB Journal, 24, 3562–3571.

8. John, G. B., et al. (2011). Role of Klotho in Aging, Phosphate Metabolism, and CKD. American Journal of Kidney Diseases, 58(1), 127–134.

9. Morishita, K., et al. (2001). The progression of aging in klotho mutant mice can be modified by dietary phosphorus and zinc. Journal of Nutrition, 131, 3182–3188.

10. Kuro-o, M. (2010). A potential link between phosphate and aging—Lessons from Klotho-deficient mice. Mechanisms of Ageing and Development, 131, 270–275.

11. Razzaque, M. S. (2011). Phosphate toxicity: new insights into an old problem. Clinical Science (Lond.), 120(3), 91–97.

12. Kuro-o, M. (2009). Klotho and aging. Biochimica et Biophysica Acta, 1790(10), 1049–1058.

13. Jacob, H. J. (2010). The Rat: a model used in biomedical research. Methods in Molecular Biology, 597, 1–11.

14. Szpirer, C. (2020). Rat models of human diseases and related phenotypes: a systematic inventory of the causative genes. Journal of Biomedical Science, 27, 84.

15. Chenouard, V., et al. (2021). Advances in Genome Editing and Application to the Generation of Genetically Modified Rat Models. Frontiers in Genetics, 12:615491.

16. Shao, Y., et al. (2014). CRISPR/Cas-mediated genome editing in the rat via direct injection of one-cell embryos. Nature Protocols, 9, 2493–2512.

17. Yoshimi, K., et al. (2014). Allele-specific genome editing and correction of disease-associated phenotypes in rats using the CRISPR-Cas platform. NatureCommunications, 5:4240.

18. Sato, M., et al. (2022). Recent Advances in the Production of Genome-Edited Rats. International Journal of Molecular Sciences, 23, 2548.

19. Cornelissen, G. (2014). Cosinor-based rhythmometry. Theoretical Biology and Medical Modelling, 11, 16.

20. Nagao, Y., Endo, H., Sakashita, E., Ohmori, T., and Hayakawa, M. (2021). Temporary treatment medium, treatment kit, embryogenesis arrest inhibitor, embryogenesis arrest inhibitory method, developmental engineering product preparation method, transplantation method, therapeutic method, and developmental engineering product. International Publication No. WO2021193596A1; Japanese Patent No. 7722712 (JP7722712B2).

